# Uncovering the dark transcriptome in polarized neuronal compartments with mcDETECT

**DOI:** 10.1101/2025.03.27.645744

**Authors:** Chenyang Yuan, Krupa Patel, Hongshun Shi, Hsiao-Lin V. Wang, Feng Wang, Ronghua Li, Yangping Li, Victor G. Corces, Hailing Shi, Sulagna Das, Jindan Yu, Peng Jin, Bing Yao, Jian Hu

**Affiliations:** Department of Human Genetics, Emory University School of Medicine, Atlanta, GA 30322, USA; Department of Biostatistics and Bioinformatics, Rollins School of Public Health, Emory University, Atlanta, GA 30322, USA; Department of Urology, Emory University School of Medicine, Atlanta, GA 30322, USA; Winship Cancer Institute, Emory University School of Medicine, Atlanta, GA 30322, USA; Emory Center for Neurodegenerative Diseases, Emory University School of Medicine, Atlanta, GA 30322, USA; Department of Cell Biology, Emory University School of Medicine, Atlanta, GA 30322, USA

**Author notes:** **Correspondence:** Bing Yao, Jian Hu. Deceased (Yangping Li).

**Keywords:** spatial transcriptomics, dark transcriptome, neuronal compartments, Alzheimer’s disease, machine learning

## Abstract

Spatial transcriptomics (ST) is a powerful tool for studying the molecular basis of brain diseases. However, most current analyses focus only on nuclear or somatic expression, overlooking distally localized mRNAs, known as the “dark transcriptome.” These transcripts, which make up nearly 40% of the brain transcriptome, are packaged into RNA granules and transported to polarized neuronal structures critical for neuronal function. Here, we present mcDETECT, a machine learning framework that leverages *in situ* ST (iST) data to detect RNA granules and resolve their RNA composition. Applying mcDETECT to simulated and real datasets from multiple iST platforms, we demonstrate robust granule detection and molecular subtyping. Granule-informed downstream analysis further revealed functionally distinct neuronal substates not captured by somatic expression alone. In an Alzheimer’s disease (AD) mouse model, mcDETECT uncovered alterations in RNA granule distributions and associated neuronal substates prior to measurable neuronal loss, highlighting potential therapeutic targets for early AD pathology.

## Introduction

Neurons are highly polarized cells with distal subcellular compartments—such as synapses, dendrites, and axons—that are essential for neural connectivity and communication. The discovery of messenger RNAs (mRNAs) localized near these compartments suggests that they achieve local autonomy by synthesizing key proteins locally to support plasticity and function^1,2^. Conventional single-cell or single-nucleus RNA sequencing (sc/snRNA-seq) techniques fail to capture these spatially localized mRNAs because they are restricted to profiling transcripts within or close to the nucleus, leaving the mRNA pools in distal compartments undetected. To highlight this critical knowledge gap, Seth A. Ament and Alexandros Poulopoulos proposed the concept of the “dark transcriptome”—a term inspired by astrophysics’ “dark matter,” which refers to mass that cannot be directly observed but exerts measurable effects^3^. The dark transcriptome represents a hidden layer of transcriptional diversity that operates at a resolution beyond cell types or subtypes, offering molecular insights complementary to traditional sequencing approaches. Remarkably, this unexplored transcriptional pool is estimated to constitute ∼40% of the total transcriptome in the adult mouse brain^4,5^, as demonstrated by both iST and fluorescence imaging of individually cultured neurons^6,7^. These compartmentalized mRNAs are transcribed in the nucleus, assembled into RNA granules by RNA-binding proteins (RBP), and then transported to distal regions^8^. Despite extensive research on RNA granules in neurons, studies have predominantly focused on their RBP components^9,10^, which are more amenable to tagging and detection. In contrast, the mRNA composition of these granules remains poorly characterized.

Recent advancements in spatial transcriptomics (ST) have enabled gene expression profiling with location information and transformed our understanding of tissue functional organization^11-16^. In particular, *in situ* hybridization-based technologies (iST), such as STARmap^17^, seqFISH^18-20^, and MERFISH^21,22^, offer gene expression measurement at subcellular resolution. Building on these techniques, methods such as Basyor^5^, JSTA^23^, FICTURE^24^, Bento^25^ and ELLA^26^ have recently been developed to analyze gene expression at the subcellular resolution. However, these methods primarily focus on mRNAs within the cell body, inferred from nuclear and/or membrane staining. As a result, distal RNA granules—which are not visible in these staining images—remain undetectable, leaving the dark transcriptome largely unexplored.

To fill this gap, we developed mcDETECT (**m**ulti-**c**hannel **DE**nsity-based spatial **T**ranscriptom**E C**lus**T**ering), a machine learning framework designed to study the dark transcriptome related to polarized compartment in brain using iST data. mcDETECT first employs density-based clustering to pinpoint the locations of distal RNA granules in three-dimensional (3D) space. Subsequently, it assigns surrounding mRNA molecules to each granule to reconstruct their transcriptome profiles. We applied mcDETECT to both simulated and real mouse brain datasets generated using three state-of-the-art iST platforms with different gene coverages. mcDETECT successfully identified multiple classes of RNA granules, each associated with distinct functional roles across brain regions. Importantly, these spatially resolved RNA granules reveal transcriptomic signatures of neuronal substates that are distinct from and complementary to those defined by somatic transcriptomes. To further demonstrate mcDETECT’s utility in disease studies, we generated iST data with a customized panel enriched for synaptic, dendritic, and axonal genes in an Alzheimer’s disease (AD) mouse model. Using this dataset, mcDETECT identified changes in the abundance and molecular composition of RNA granules as indicators of early AD. These granule-level alterations also revealed shifts in neuronal substates that precede detectable changes in neuronal somata. Overall, mcDETECT opens a new avenue for the neuroscience community to profile the dark transcriptome, uncover molecular insights, and identify potential RNA granule-level targets for therapeutic intervention in neurodegenerative diseases.

## Results

### Overview of mcDETECT

The workflow of mcDETECT is illustrated in **Fig. 1**. It begins by examining the subcellular distribution of mRNAs in an iST sample. Each mRNA molecule is treated as a distinct point with its own 3D spatial coordinates considering the thickness of the sample. Unlike many cell-type marker genes, which are typically found within the nucleus or soma, compartmentalized mRNAs often form small aggregates outside the soma (**Fig. 1a**). mcDETECT uses a density-based clustering^27^ approach detailed in **Fig. 1b** to identify these extrasomatic aggregates. This involves calculating the Euclidean distance between mRNA points and defining the neighborhood of each point within a specified search radius. Points are then categorized as core points, border points, or noise points based on their reachability from neighboring points. mcDETECT recognizes each connected bundle of core and border points as an mRNA aggregate. To minimize false positives, it excludes aggregates that substantially overlap with somata, which are estimated by dilating the nuclear masks derived from DAPI staining. mcDETECT then repeats this process for multiple granule markers, merging aggregates from different markers that exhibit high spatial overlap (**Fig. 1c**). After aggregating across all markers, an additional filtering step removes aggregates containing mRNAs from negative control genes, which are known to be enriched exclusively in nuclei and somata. The remaining aggregates are considered individual RNA granules. As shown in **Fig. 1d**, mcDETECT then computes the minimum enclosing sphere for each aggregate to connect neighboring mRNA molecules from all measured genes and summarizes their counts, thereby defining the spatial transcriptome profile of individual RNA granules. mcDETECT also supports downstream applications, as illustrated in **Fig. 1e**. For example, it can classify RNA granules into molecular subtypes and analyze their spatial relationships with neurons. This approach enables the identification of neuronal substates based on the types of neighboring granules—information that cannot be resolved using somatic transcriptomes alone. By comparing disease and control tissues, mcDETECT can also identify granule-associated biomarkers, such as alterations in granule density linked to neurological disorders.

**Fig. 1.**
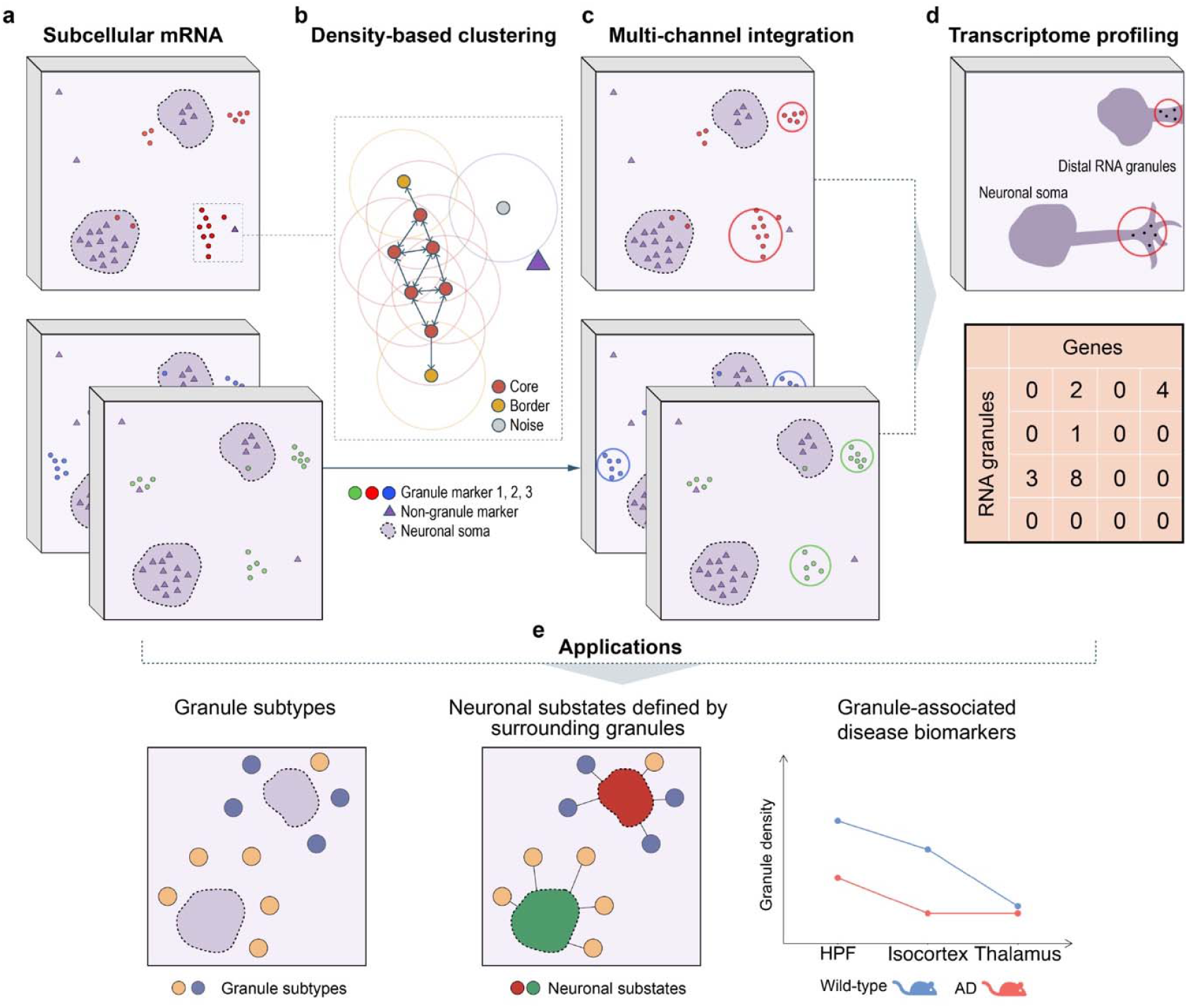
Workflow of mcDETECT. **a**. Schematic representation of the subcellular spatial distribution of compartmentalized mRNAs (red, blue, and green dots) and non-compartmentalized mRNAs (purple triangles) within neurons. **b**. mcDETECT applies a density-based clustering approach to identify extrasomatic mRNA aggregates for each distally localized gene. **c**. mcDETECT repeats this process for all distally localized genes, merges aggregates with significant spatial overlap, and identifies individual RNA granule locations. **d**. mcDETECT collects all mRNAs within the minimal enclosing sphere of each aggregate to construct the transcriptome profile of individual RNA granules. **e**. Example downstream applications of mcDETECT, including the identification of granule subtypes, granule-informed neuronal substates, and granule-associated disease biomarkers.

### Selection of markers for distal RNA granules

The first step in mcDETECT is to select a set of marker genes associated with distal RNA granules. RNA granules are a diverse family and can be classified by their protein composition into stress granules, processing bodies, and transport granules. In most tissues, RNA granules represent only a small fraction of the overall transcriptome. The brain, however, presents a notable exception. Due to the highly polarized architecture of neurons, transport granules constitute a much large portion (∼40%) of neuronal transcriptome^3-5^. This reflects their critical role in delivering mRNAs to distal neuronal compartments, such as synapses, dendrites, and axons, to support local protein synthesis. To capture these RNA granules, we compiled a list of markers associated with synaptic, dendritic, and axonal structures and functions. As detailed in **Supplementary Table 2**, these markers have been reported in multiple studies to be enriched in distal neuronal compartments, supporting their packaging into RNA granules for transport. To validate the subcellular enrichment patterns of our selected genes, we analyzed their RNA localization in brain datasets generated by three different iST platforms. As shown in **Fig. 2a**, across all three platforms, our selected genes exhibit significantly higher extrasomatic localization compared to other genes (one-sided t-test: p = 9.1 × 10^-6^, 0.0013, 0.0021 for Xenium 5K, MERSCOPE, and CosMx), confirming their preferential distribution to distal compartments. To further assess the functional relevance of these distally localized mRNAs, we analyzed their ribosome association using a mouse coronal hemisphere dataset generated using RIBOmap, a highly multiplexed 3D *in situ* technique for translatome profiling^28^. For each gene, we quantified the relative proportion of ribosome-bound mRNAs in peripheral neuronal processes versus somata. Notably, the 26 genes that overlap with our target set demonstrate significantly greater ribosome association in peripheral compartments (median = 22.1%) compared to the remaining 5,387 genes (median = 18.3%, p = 0.00024; **Fig. 2b**). These results suggest that these distal mRNAs are actively translated in neuronal processes, supporting their functional role in shaping neuronal connectivity.

**Fig. 2.**
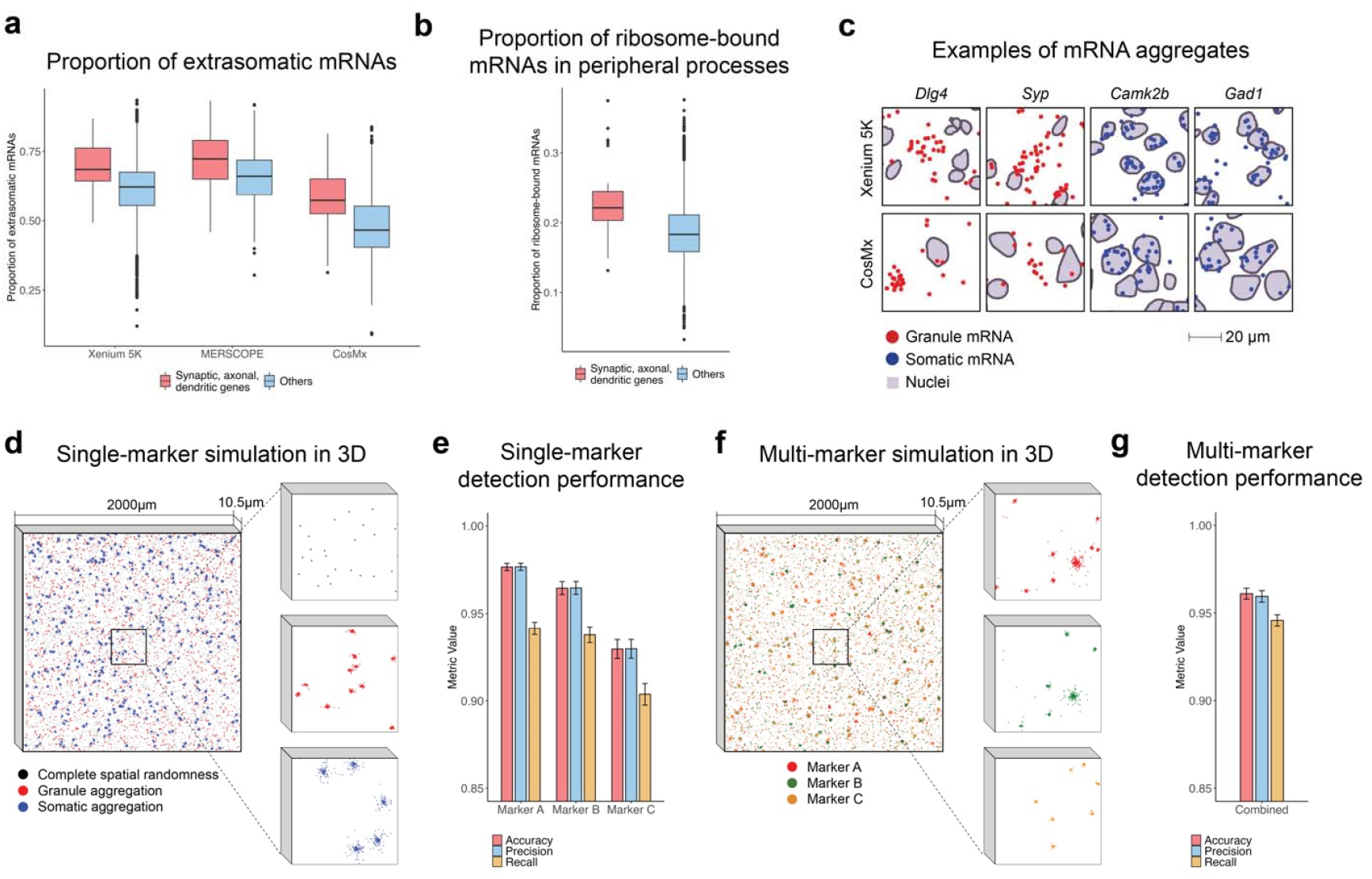
Granule marker selection and simulation study. **a**. Boxplot comparing the proportion of extrasomatic mRNAs for selected synaptic, dendritic, and axonal genes versus other genes in the Xenium 5K, MERSCOPE, and CosMx datasets. The lower and upper hinges correspond to the first and third quartiles, and the center represents the median. The upper (lower) whisker extends from the hinge to the largest (smallest) value within 1.5 × the interquartile range. Data beyond the whiskers are plotted individually. **b**. Boxplot comparing the proportion of ribosome-bound mRNAs in peripheral neuronal processes for selected synaptic, dendritic, and axonal genes versus other genes in the RIBOmap dataset. Boxplot definitions are the same as in **a. c**. Examples of subcellular mRNA aggregation patterns for synaptic markers (*Dlg4, Syp*) and neuronal markers (*Camk2b, Gad1*) in the Xenium 5K and CosMx datasets. Synaptic and neuronal mRNAs are shown as red and blue dots, respectively. Neuronal nuclei are identified from DAPI staining and highlighted in purple. Scale bar: 20 μm. **d**. Left: Simulation of a single granule marker in 3D space, where each dot represents a subcellular mRNA molecule. Black, red, and blue dots indicate complete spatial randomness (CSR), aggregation within distal compartments, and aggregation within somata, respectively. Right: Zoomed-in view of the three distribution patterns in a small region. **e**. Bar plot showing the detection accuracy, precision, and recall achieved by mcDETECT on the simulated 3D single-marker data across 200 simulation runs. Bar heights represent mean values, and error bars denote mean ± one standard deviation. **f**. Left: Simulation of three granule markers in 3D space, where each dot represents a subcellular mRNA molecule. Red, green, and orange dots indicate marker types A, B, and C, respectively. Right: Zoomed-in view of a small region stratified by marker type. **g**. Bar plot showing the detection accuracy, precision, and recall achieved by mcDETECT on the simulated 3D multi-marker data across 200 simulation runs. Bar plot definitions are the same as in **e**.

### Simulation study

To evaluate the effectiveness of mcDETECT in identifying RNA granules, we conducted a simulation study to benchmark its performance. We simulated a 2000 × 2000 μm^2^ tissue area with a thickness 10.5 μm, mimicking the iST sample setting. In real iST data, we find that synaptic markers such as *Dlg4* and *Syp* consistently exhibit clear extrasomatic aggregation across both platforms, while neuronal markers *Camk2b* and *Gad1* are predominantly enriched in the nuclear region (**Fig. 2c**). Inspired by these mRNA distribution patterns in real data^29^, we simulated three distally localized genes (denoted A, B, and C) with varying overall densities. Each gene encompassed three distinct subcellular mRNA distribution patterns, as illustrated in **Fig. 2d:** (1) Complete spatial randomness (CSR), where mRNA molecules are randomly distributed without forming aggregates. (2) Aggregation within distal compartments, modeled by simulating mRNA clusters with a mean radius of 1 μm, reflecting the reported granule size in existing studies^8-10,29^. We randomly set 80% of the points in these clusters to be located outside the somata, with the remaining 20% distributed within somata. (3) Aggregation within somata, modeled by simulating mRNA clusters with a mean radius of 3.5 μm, with 80% of the points located within somata. For each gene, we conducted 200 simulation runs and applied mcDETECT to detect the distal RNA granules. As illustrated in the bar plots in **Fig. 2e**, mcDETECT achieved overall accuracies of 97.7%, 96.4%, and 93.0% for markers A, B, and C, respectively. Notably, mcDETECT was not only sensitive to true distal RNA granules—reflected by mean recalls of 94.1%, 93.8%, and 90.4%—but also effectively filtered out most intrasomatic aggregates, as indicated by mean precisions of 97.7%, 96.5%, and 93.0% for markers A, B, and C, respectively.

The above simulation focused on RNA granule detection using a single marker. In scenarios involving multiple granule markers, mcDETECT enhances statistical power by integrating aggregations from multiple markers while accounting for their colocalization patterns. To demonstrate this capability, we simulated a scenario with three distally localized genes, A, B, and C, whose mRNA molecules were visualized in red, green, and yellow in **Fig. 2f**. These three markers partially co-clustered within the same 3D space, and using any single marker recovered only a subset of the total RNA granules, with a recall of approximately 60%. In contrast, when mcDETECT combined all three markers, it achieved an accuracy of 96.1%, a mean precision of 96.0%, and a mean recall of 94.6% across 200 simulation runs, as shown in the bar plot in **Fig. 2g**.

In most iST data analyses, the x–y plane typically spans a much larger area (e.g., 10 × 10 mm^2^) than the z-axis (∼10 μm). As a result, existing approaches often treat tissues as two-dimensional (2D) sections and ignore their thickness. By conducting additional simulations in a 2D setting, we demonstrate that omitting the z-axis can lead to false-positive RNA granule detection. This occurs because RNA projections are condensed into a 2D plane, making them appear aggregated when they are not in 3D. (**Supplementary Note 1**).

### Application to mouse brain 10x Xenium 5K data

To demonstrate the capabilities of mcDETECT on real data, we applied it to a dataset from a 9-week-old wild-type (WT) mouse brain, generated using the 10x Xenium 5K platform^30^. This dataset has a broad coverage of 5,006 genes, including 24 of our selected granule markers. Using these markers, mcDETECT identified 170,501 RNA granules across ten brain regions. As shown in **Fig. 3a**, these granules are more concentrated in the cortical layers, hippocampus (HPF), and olfactory areas (OLF). Directly evaluating the accuracy of granule detection in real data is challenging due to technical limitations in obtaining ground truth on the same iST slide. Unlike cells, granules are difficult to capture using protein staining due to their complex protein composition and lack of specific antibody markers. Therefore, we evaluated mcDETECT’s output in two aspects. First, we examined the sizes of the identified granules. As shown in **Fig. 3b**, the radii of 90% of the identified granules fall within the range of 0.54–1.69 μm, with a median of 0.93 μm, which is consistent with previously reported granule sizes^31-33^. Second, we compared the transcriptomic profiles of the identified granules with those of neuronal somata. **Fig. 3c** presents a heatmap of 10 granule markers and 10 neuronal markers^34-37^ used as negative controls. In the identified granules, granule markers are highly enriched, whereas negative control mRNAs are sparsely detected. In contrast, neuronal somata exhibit high expression of both granule and neuronal markers. This distinct enrichment pattern confirms that the identified granules are not soma fragments or random cellular debris.

**Fig. 3.**
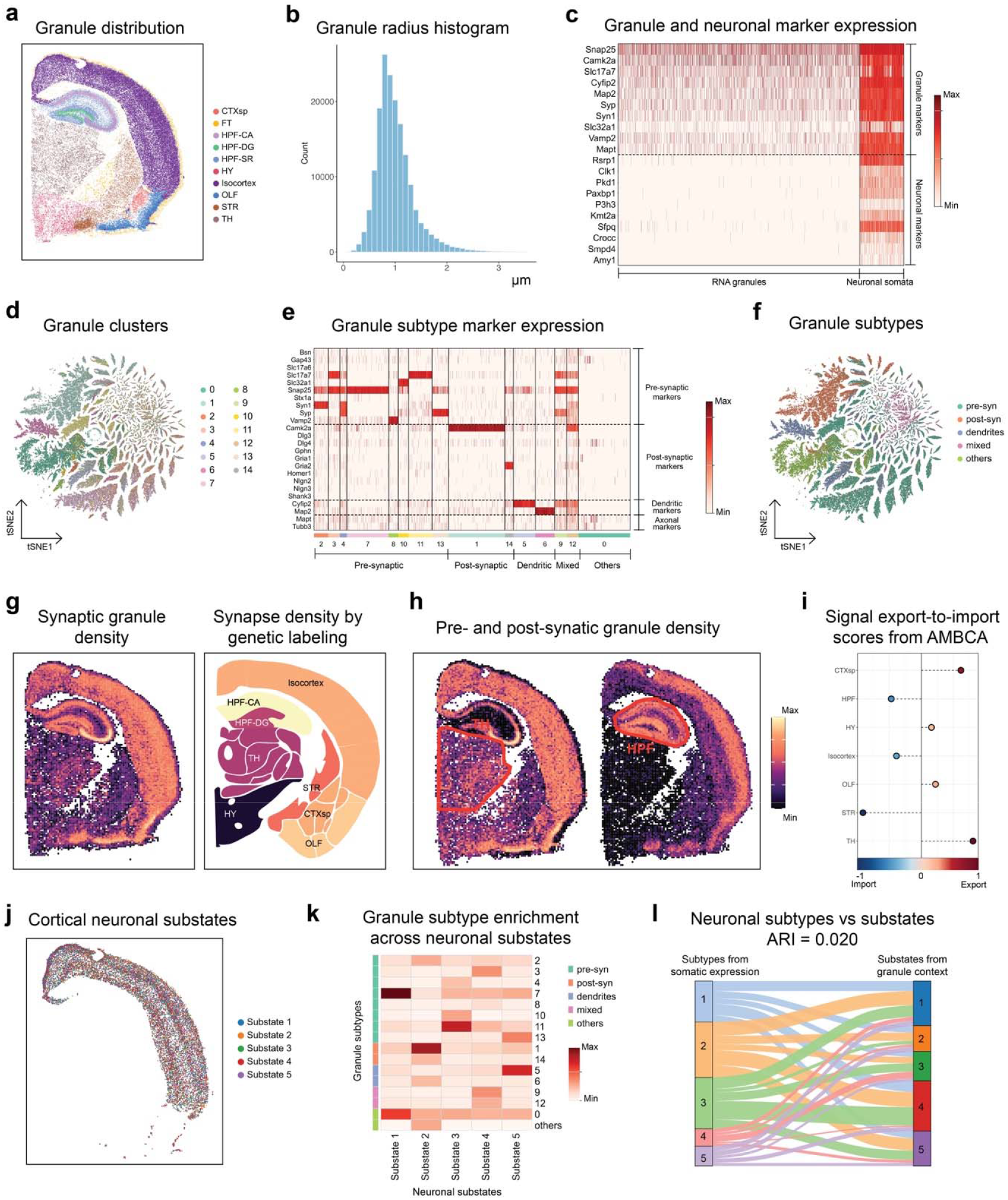
Application to the Xenium 5K dataset. **a**. Scatter plot showing all RNA granules identified by mcDETECT across ten major brain regions: cortical subplate (CTXsp), fiber tracts (FT), cornu ammonis of the hippocampus (HPF-CA), dentate gyrus of the hippocampus (HPF-DG), stratum radiatum of the hippocampus (HPF-SR), hypothalamus (HY), isocortex, olfactory areas (OLF), striatum (STR), and thalamus (TH). **b**. Histogram showing the distribution of radii for all identified RNA granules. **c**. Heatmap showing the enrichment patterns of 10 granule markers and 10 neuronal markers in RNA granules and neuronal somata. Color represents the standardized enrichment level of each gene within each group. **d**. t-SNE plot of all RNA granules, colored by 15 clusters obtained from K-Means clustering based on enrichment of 24 granule-subtype-specific genes (pre-synaptic, post-synaptic, dendritic, and axonal). **e**. Heatmap showing the enrichment patterns of the 24 granule-subtype-specific genes across the 15 granule clusters. Color definitions are the same as in **c. f**. t-SNE plot of all RNA granules, colored by five major granule subtypes defined by dominant subtype marker enrichment. **g**. Left: scatter plot showing the number of synaptic granules (i.e., pre- or post-synaptic) identified by mcDETECT in 50 × 50 μm^2^ spots spanning the tissue slice. Right: regional synapse densities estimated using volume electron microscopy and genetic labeling on a coronal slice corresponding to the approximate bregma position in **a. h**. Scatter plots showing the number of pre- and post-synaptic granules in each 50 × 50 μm^2^ spot. The TH and HPF regions are manually annotated. **i**. Dot plot summarizing the signal export-to-import score for each brain region, with regions colored according to whether they predominantly receive (blue) or send (red) signals, based on estimated pairwise axonal projection strengths from the Allen Mouse Brain Connectivity Atlas. **j**. Scatter plot showing the five cortical neuronal substates defined based on granule context. **k**. Heatmap showing the enrichment scores of granule clusters and their associated subtypes within each neuronal substate, calculated by averaging the granule context embeddings of individual neurons. **l**. Alluvial plot illustrating the correspondence between neuronal subtypes derived from somatic transcriptome clustering (left; K-Means clustering) and neuronal substates defined by granule context (right).

After recovering granule locations, mcDETECT enables the construction of transcriptome profiles for individual RNA granules, facilitating subtype classification and functional analysis. The identified granules contain a median of 16 mRNA molecules per granule, with 72 genes detected in more than 5% of all granules. To study their molecular subtypes, we performed K-Means clustering based on granule-level gene enrichment and identified 15 initial clusters, visualized by t-SNE in **Fig. 3d**. Based on heatmap of selected markers in **Fig. 3e**, we further annotated the granules as enriched for pre-synaptic genes, post-synaptic genes, dendritic genes, or mixed types in **Fig. 3f**. We next demonstrate the biological significance of these granule subtypes in three aspects. First, granule subtypes can serve as quantitative biomarkers of its associated functional compartments. Specifically, we calculated the density of synapse-associated granules (enriched for both pre- and post-synaptic genes) within each 50 × 50 μm^2^ spot across ten major brain regions (**Fig. 3g, left**). Notably, these granule densities showed a strong correlation with synapse densities derived from age-matched WT mouse brains using volume electron microscopy and genetic labeling^38^ (**Fig. 3g, right**), with a weighted Spearman correlation coefficient of 0.98. This finding provides orthogonal validation of the accuracy of our granule detection method and underscores its biological significance as a proxy for synapse abundance.

Second, granule subtypes provide a dynamic readout of cellular function. Synapses are functionally divided into two major compartments: the pre-synapse, which releases neurotransmitters to initiate signal transmission, and the post-synapse, which contains receptors that detect and respond to these signals. The balance between pre- and post-synapses is a critical indicator of signal flow directionality and regional specialization within the brain. However, examining this balance is challenging, as current techniques rely on protein antibodies to isolate these subcellular compartments but cannot preserve their native spatial context. Using mcDETECT, we overcame this limitation by shifting from directly identifying compartments to quantifying the distributions of their associated granules. As shown in **Fig. 3h**, the thalamus (TH) contains a higher concentration of pre-synaptic granules, indicating its enriched activities related to signal export. In contrast, the HPF exhibits a greater density of postsynaptic granules, indicating its enriched activities related to signal reception and processing. To validate this finding, we compared our results with data from the Allen Mouse Brain Connectivity Atlas (AMBCA), which maps axonal projection strengths by tracking the trajectories of enhanced green fluorescent protein (EGFP)-expressing adeno-associated viral vectors. By summarizing inward and outward projections for each region, we computed a signal export-to-import score, as detailed in **Fig. 3i**. The TH exhibits a predominance of outgoing projections, consistent with its role as an output hub and its enrichment of pre-synaptic granules. The HPF, on the other hand, receives more incoming projections and is enriched with dendritic spines, accounting for its high abundance of post-synaptic granules.

Third, the spatial organization of granules around neurons reveals neuronal substates that cannot be resolved by their somatic transcriptomes alone. As a proof of concept, we focused our analysis on the isocortex, given its high neuronal density and functional significance in cognition. For each neuron, we identified its surrounding granules within a predefined distance. We then generated a granule context embedding that incorporated both granule subtypes and their relative distances to the neuronal somata. Latent Dirichlet Allocation (LDA) clustering of these embeddings revealed five neuronal substates in the isocortex, denoted as Substate 1–5, each exhibiting distinct laminar distributions across cortical layers (**Fig. 3j**). These neuronal substates are characterized by distinct spatial compositions of granule subtypes in their local microenvironment (**Fig. 3k**). Notably, they could not be resolved by somatic transcriptome clustering alone, as demonstrated in the Alluvial plot in **Fig. 3l**, which shows little correspondence with neuronal subtypes derived from highly expressed somatic genes and yields a low Adjusted Rand Index (ARI) of 0.02. We hypothesized that these substates, defined by extrasomatic RNA granules, reflect neurons’ specialized functional roles and would be partially mirrored in their somatic gene expression. To test this, we performed differential expression (DE) and Gene Ontology (GO) analyses on the somatic transcriptomes of each substate, identifying distinct marker genes and significantly upregulated pathways. For example, Substate 1 neurons, surrounded by abundant pre-synaptic granules, showed upregulation of pathways related to pre-synaptic membrane potential, neurotransmitter uptake, and L-glutamate transport, suggesting a role in excitatory transmission and synaptic output. In contrast, Substate 2 neurons exhibited enrichment of post-synaptic granules and elevated expression of genes involved in synaptic plasticity, long-term potentiation (LTP), and dendritic spine development, indicating their specialization in synaptic input and experience-dependent remodeling. Supporting figure panels for Substates 1 and 2 are provided in **Supplementary Note 2**, while a full interpretation of the remaining substates is given in **Supplementary Note 3**. These results demonstrate that granule-defined substates represent functionally distinct neuronal populations.

### Comparing WT and AD mouse brains using MERSCOPE data with a customized gene panel

We next demonstrate mcDETECT’s ability to uncover disease-relevant insights through granule-level analysis. This task is challenging with existing iST data due to either limited granule marker coverage or shallow sequencing depth in high-coverage platforms such as Xenium 5K. To address such limitations, we designed a 290-gene panel tailored for profiling distal RNA granules (**Supplementary Table 5**), comprising 48 synaptic markers, 12 dendritic markers, and 9 axonal markers. The remaining genes include cell type and brain region markers, ensuring the panel remains compatible with standard analyses. Using this customized panel, we performed MERSCOPE iST experiments on coronal brain sections from two 8-week-old mice: one C57BL/6J (WT model) and one 5xFAD (AD model). This time point corresponds to the pre-symptomatic stage of AD, when minimal neuronal degeneration is typically observed. Using the granule markers, mcDETECT identified a total of 1,498,704 RNA granules in the WT and AD samples (**Fig. 4a**). Compared to the Xenium 5K dataset, the higher granule count in MERSCOPE can be attributed to differences in tissue area, gene coverage, and platform-specific mRNA capture efficiency (**Supplementary Note 4**). To compare the WT and AD samples, we first evaluated potential granule-level batch effects. As shown in **Fig. 4b**, the granule size distributions were similar between samples, with median radii of 0.95 μm in WT and 0.96 μm in AD. Furthermore, t-SNE visualization based on gene enrichment revealed substantial intermixing of granules from both groups (**Fig. 4c**), indicating minimal batch effects and supporting subsequent joint analysis. Following the same analysis pipeline, we identified 15 initial granule clusters and annotated each as being enriched for pre-synaptic genes, post-synaptic genes, dendritic genes, or mixed types (**Fig. 4d–f**). Similar to the Xenium 5K data, the synaptic granules in the MERSCOPE WT sample also showed a strong correlation with synapse density (**Supplementary Note 5**).

**Fig. 4.**
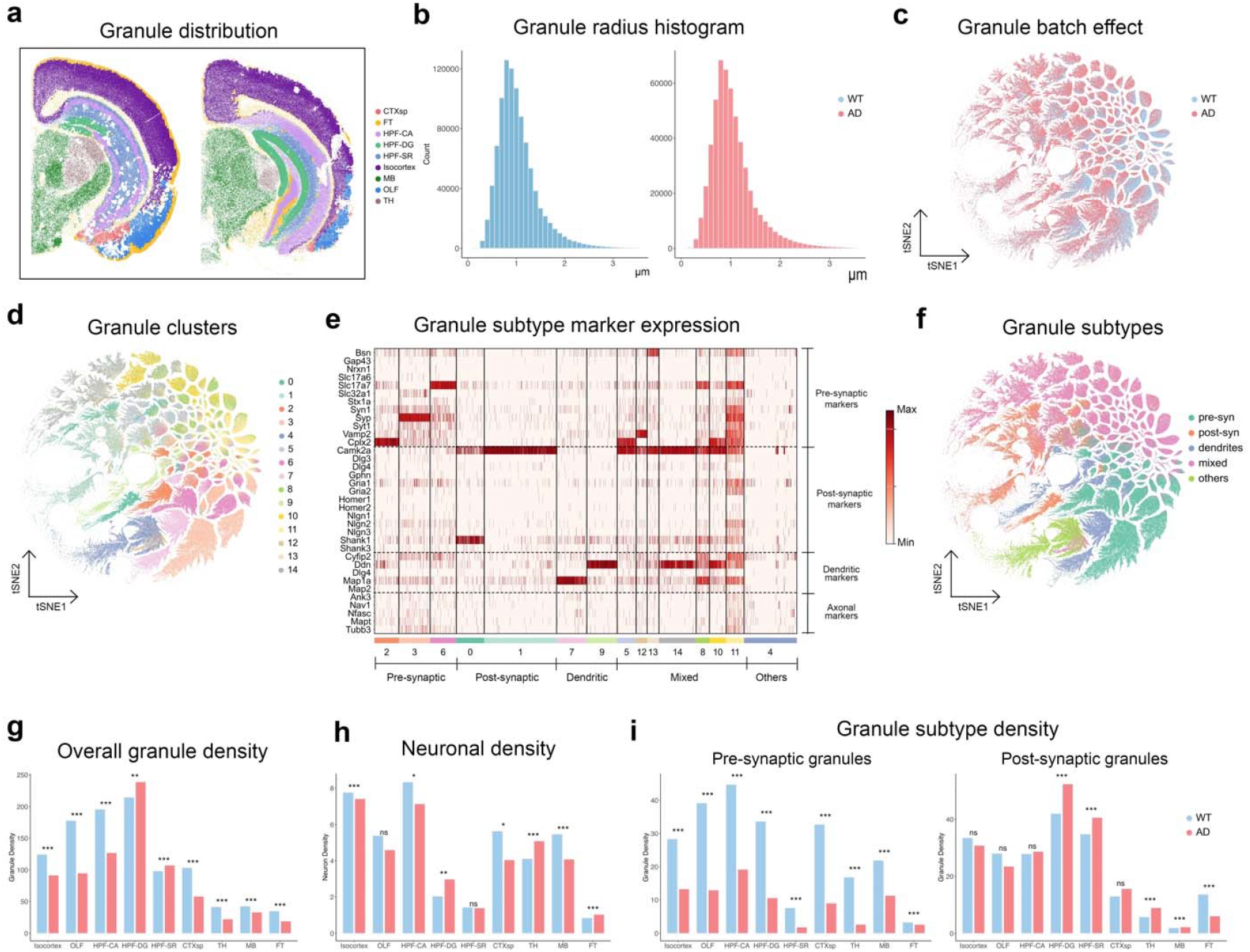
Granule alterations between wild-type (WT) and Alzheimer’s disease (AD) mouse brains in MERSCOPE datasets. **a**. Scatter plot showing all RNA granules identified by mcDETECT in WT and AD samples across nine major brain regions: cortical subplate (CTXsp), fiber tracts (FT), cornu ammonis of the hippocampus (HPF-CA), dentate gyrus of the hippocampus (HPF-DG), stratum radiatum of the hippocampus (HPF-SR), isocortex, midbrain (MB), olfactory areas (OLF), and thalamus (TH). **b**. Density plots comparing the distribution of radii for all identified RNA granules between WT and AD samples. **c**. t-SNE plot of all RNA granules, colored by sample identity (WT vs AD). **d**. t-SNE plot of all RNA granules, colored by 15 clusters obtained from K-Means clustering based on enrichment of 34 granule-subtype-specific genes (pre-synaptic, post-synaptic, dendritic, and axonal). **e**. Heatmap showing the enrichment patterns of the 34 granule-subtype-specific genes across the 15 granule clusters. Color represents the standardized enrichment level of each gene within each group. **f**. t-SNE plot of all RNA granules, colored by five major granule subtypes defined by dominant subtype marker enrichment. **g**. Grouped bar plot comparing regional RNA granule densities (measured as the number of granules per 50 × 50 μm^2^ spot) between WT and AD samples, after adjusting for differences in capture efficiency. Two-sample t-tests were performed on log-transformed spot-level granule counts within each brain region. ns: not significant; *: p < 0.05; **: p < 0.01; ***: p < 0.001. **h**. Grouped bar plot comparing regional neuronal densities between WT and AD samples. Two-sample t-tests were performed on log-transformed spot-level neuron counts within each brain region. ns: not significant; *: p < 0.05; **: p < 0.01; ***: p < 0.001. **i**. Grouped bar plots comparing regional densities of pre- and post-synaptic granules between WT and AD samples, after adjusting for differences in capture efficiency. Two-sample t-tests were performed on log-transformed spot-level granule counts within each brain region. ns: not significant; *: p < 0.05; **: p < 0.01; ***: p < 0.001.

We subsequently compared the distribution of RNA granules and their subtypes between WT and AD samples. As shown in **Fig. 4g**, the overall granule density was significantly reduced in the AD sample compared to WT, even after adjusting for differences in mRNA capture efficiency between the two experiments. This reduction was consistent across multiple brain regions, most notably in the isocortex, OLF, and cornu ammonis (HPF-CA). Neuronal densities within each brain region, however, remained largely unchanged (**Fig. 4h**). This discrepancy suggests that disruption of RNA granule transport may serve as an early and detectable hallmark of AD pathology, potentially preceding measurable neuronal loss. We further examined changes in granule subtype densities. As shown in **Fig. 4i (left)**, the density of pre-synaptic granules was significantly reduced across all brain regions, with the most severe loss in the isocortex, OLF, and HPF-CA, regions previously reported to be most affected in early AD pathology^39,40^. In contrast, post-synaptic granule density remained relatively stable, with no significant changes in these regions (**Fig. 4i, right**). This differential response in RNA granules highlights the selective vulnerability of the pre-synapse to early AD pathology and is consistent with previous studies showing significant pre-synapse loss, whereas post-synapse density remains relatively preserved, as observed in the J20 AD mouse model^41^. Other granule subtypes, including those enriched for dendritic genes and mixed signals, also exhibited region-specific alterations in AD (**Supplementary Note 6**).

Next, we investigated whether any granule-defined neuronal substates exhibited disease-associated alterations. Using a similar pipeline, we clustered granule context embeddings and identified five cortical neuronal substates (Substates 1–5) that were shared between WT and AD samples but showed distinct spatial distribution patterns (**Fig. 5a**). Each substate was defined by a unique composition of granule subtypes within its local microenvironment, as shown in the heatmap in **Fig. 5b**. Consistent with our previous findings, the Alluvial plot in **Fig. 5c** revealed little correspondence between these context-defined substates and those derived from somatic gene expression. A low ARI of 0.026 further confirmed that granule-based substates could not be identified through somatic transcriptome analysis alone.

**Fig. 5.**
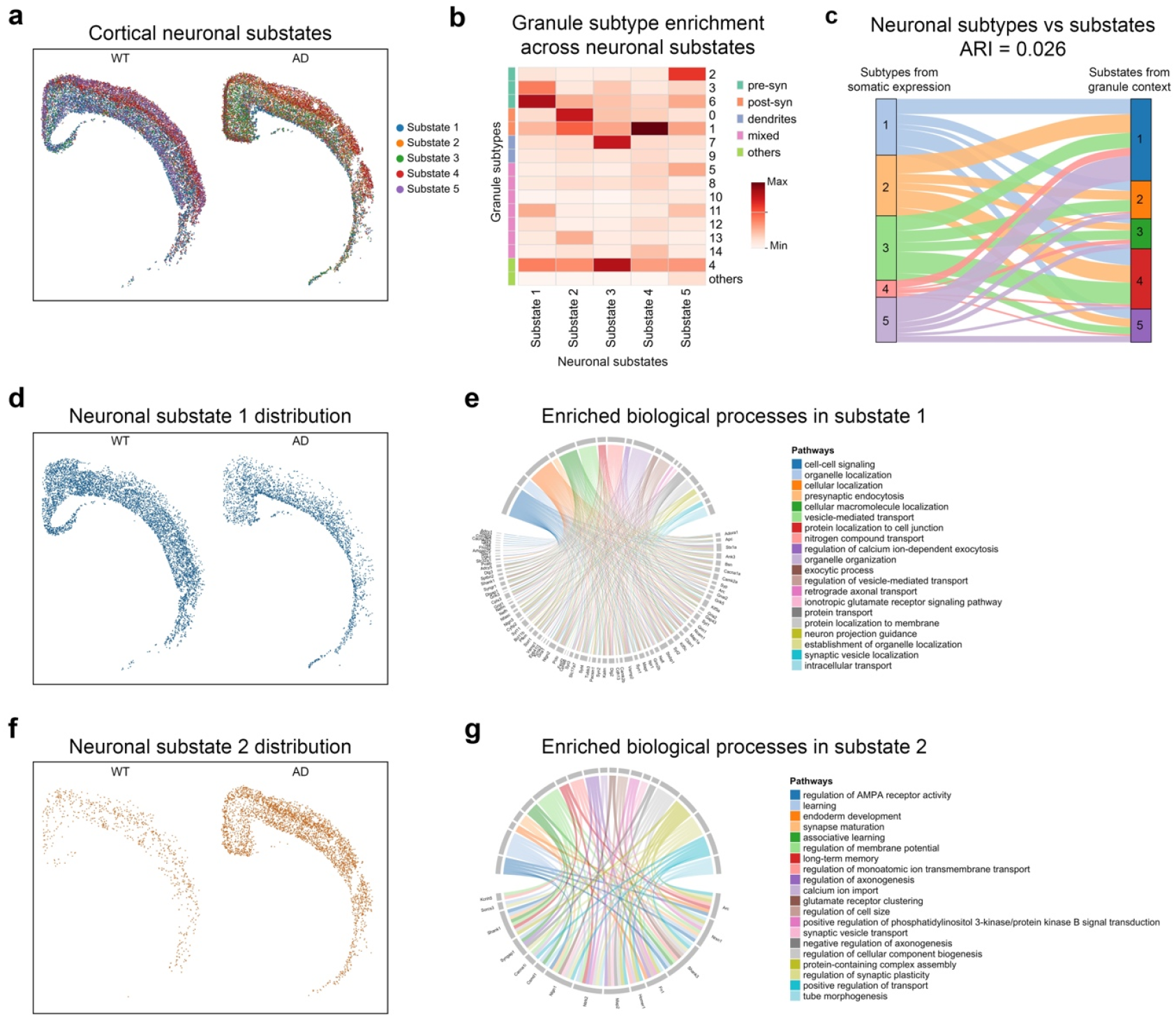
Cortical neuronal substate alterations between wild-type (WT) and Alzheimer’s disease (AD) mouse brains in MERSCOPE datasets. **a**. Scatter plot showing the five cortical neuronal substates defined based on granule context. **b**. Heatmap showing the enrichment scores of granule clusters and their associated subtypes within each neuronal substate, calculated by averaging the granule context embeddings of individual neurons. **c**. Alluvial plot illustrating the correspondence between neuronal subtypes derived from somatic transcriptome clustering (left; K-Means clustering) and neuronal substates defined by granule context (right). **d**. Scatter plot showing the distribution of Substate 1 neurons in WT and AD samples. **e**. Chord diagram illustrating enriched biological processes and their associated genes in Substate 1 neurons. Links represent gene–pathway associations based on GO enrichment analysis. **f**. Scatter plot showing the distribution of Substate 2 neurons in WT and AD samples. **g**. Chord diagram illustrating enriched biological processes and their associated genes in Substate 2 neurons. Links are defined as in **d.**

We then asked how alterations in neuronal substates reflect the dysregulation of neural networks during AD pathology. Mapping substate density revealed striking differences between WT and AD samples in the isocortex. Most notably, Substate 1 showed the strongest decrease in AD (**Fig. 5d**). This substate represents neurons specialized in precise neurotransmitter release and excitatory signal propagation, supported by the enrichment of pathways such as presynaptic endocytosis, calcium-dependent exocytosis, glutamatergic signaling, and axonal transport (**Fig. 5e**). The depletion of Substate 1 suggests an early and critical impairment of presynaptic machinery, which has been linked to the buildup of amyloid-beta oligomers and tau pathology^42^. Such a change can weaken the brain’s capacity to initiate neurotransmission, silencing key excitatory circuits and disrupting information flow. In contrast, Substate 2 showed the greatest increase in AD (**Fig. 5f**). This substate comprises neurons with enhanced postsynaptic responsiveness and plasticity, marked by the upregulation of pathways involved in AMPA receptor regulation, glutamate receptor clustering, synapse maturation, and learning-associated plasticity (**Fig. 5g**). In AD, enhanced postsynaptic responsiveness is generally associated with pathological synaptic activity, driven by factors such as amyloid-beta oligomers, which disrupt normal plasticity and lead to impaired learning and memory^43^. The expansion of Substate 2 suggests a potential compensatory postsynaptic response to diminished presynaptic input, enhancing receptor clustering and dendritic spine development to help sustain circuit activity. Together, the opposing trajectories of these substates highlight a profound dysregulation of excitatory signaling in early AD prior to noticeable neuronal loss. The decline of stable, signal-sending presynaptic neurons (Substate 1), combined with the expansion of hyperresponsive, signal-receiving postsynaptic neurons (Substate 2), drives cortical networks toward hyperexcitability—a well-established hallmark of early AD pathology observed in both human studies and multiple mouse models^44-46^.

A detailed evaluation of other cortical neuronal substates and their biological relevance is presented in **Supplementary Note 7**. Beyond the isocortex, we extended our analysis framework to hippocampal neurons, where we consistently observed a disruption of excitatory–inhibitory balance along with additional signatures of AD-induced circuit dysfunction (**Supplementary Note 8**). These findings highlight mcDETECT as a powerful tool for leveraging granule-informed microenvironments to study the neuronal vulnerability and adaption in neurodegenerative disease.

### Application to datasets with limited granule marker coverage

We have shown that mcDETECT accurately identifies distal RNA granules in the Xenium 5K and in-house MERSCOPE datasets. These datasets cover a number of granule markers, either through a broad gene coverage or a specialized panel. However, many existing brain iST datasets contain only 300–500 genes and are not specifically designed to profile distal RNA granules. To assess mcDETECT’s scalability under such constraints, we evaluated whether a focuses subset of key granule genes could recover most RNA granules. In the Xenium 5K and in-house MERSCOPE samples, we first detected granules for each marker individually and then merged the aggregates in descending order of gene expression. As shown in **Fig. 6a**, the cumulative granule count plateaued after only a small number of top-expressed markers, with the top six accounting for over 80% of detected granules in all three datasets. Benchmarking further revealed that removing top-expressed markers such as *Snap25* or *Slc17a7* had minimal impact on the spatial granule distribution, which remained highly correlated with the full-marker results (**Supplementary Note 9**). These results demonstrate that mcDETECT remains effective even with limited marker coverage, thus enabling reanalysis of such datasets to gain deeper insights into brain organization and disease mechanisms.

**Fig. 6.**
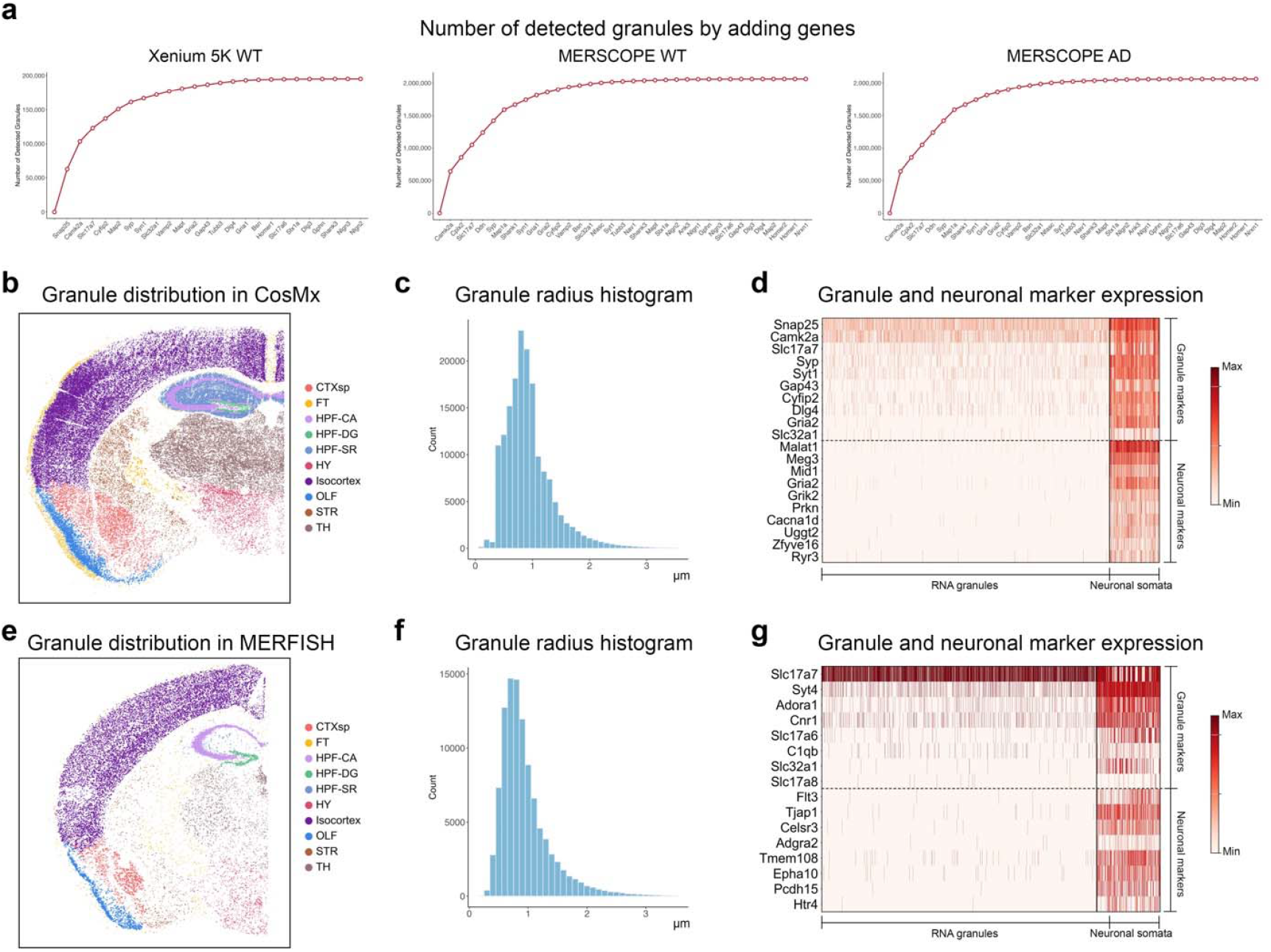
Benchmarking mcDETECT on datasets with limited granule marker coverage. **a**. Line plots showing the cumulative number of RNA granules identified by sequentially applying mcDETECT to individual granule markers and aggregating the results in the Xenium 5K, MERSCOPE WT, and MERSCOPE AD samples. **b–d**. Application to the CosMx dataset. **b**. Scatter plot showing all RNA granules identified by mcDETECT across ten major brain regions: cortical subplate (CTXsp), fiber tracts (FT), cornu ammonis of the hippocampus (HPF-CA), dentate gyrus of the hippocampus (HPF-DG), stratum radiatum of the hippocampus (HPF-SR), hypothalamus (HY), isocortex, olfactory areas (OLF), striatum (STR), and thalamus (TH). **c**. Histogram showing the distribution of radii for all identified RNA granules. **d**. Heatmap showing the enrichment patterns of 10 granule markers and 10 neuronal markers in RNA granules and neuronal somata. Color represents the standardized enrichment level of each gene within each group. **e–g**. Application to the MERFISH dataset. **e**. Scatter plot showing all RNA granules identified by mcDETECT across the same ten brain regions as in **b. f**. Histogram showing the distribution of radii for all identified RNA granules. **g**. Heatmap showing the enrichment patterns of 8 granule markers and 8 neuronal markers in RNA granules and neuronal somata. Color definitions are the same as in **d.**

To further illustrate this capability, we applied mcDETECT to two public mouse brain datasets generated using the CosMx^47^ and MERFISH^48^ platforms, each with limited coverage of canonical synaptic, dendritic, and axonal genes. Both the CosMx (**Fig. 6b–d**) and MERFISH (**Fig. 6e–g**) datasets showed results consistent with our earlier findings. Using 12 and 8 granule markers, respectively, mcDETECT identified 167,988 and 102,920 RNA granules in the CosMx and MERFISH datasets (**Fig. 6b, 6e**). The granule size distributions were similar to previous results, with median radii of 0.94 μm and 0.95 μm, respectively (**Fig. 6c, 6f**). The granule and neuronal markers also showed distinct enrichment patterns between the identified granules and neuronal somata (**Fig. 6d, 6g**), highlighting that the granules are not soma fragments or random cellular debris. We further classified the granules into molecular subtypes based on the enrichment of available subtype-specific genes, identifying populations enriched for pre- and post-synaptic markers. Notably, the spatial distributions of these synaptic granules were highly correlated with regional synapse densities from volume electron microscopy and genetic labeling, yielding weighted Spearman correlation coefficients of 0.81 and 0.90, respectively. Together, these findings highlight mcDETECT’s versatility in recovering RNA granules across diverse iST platforms, even with restricted marker panels.

## Discussion

We introduce mcDETECT, a machine learning framework designed to investigate the dark transcriptome in iST data. Whereas most existing iST analysis tools focus on nuclear or somatic transcripts, they often overlook RNA localized in distal neuronal compartments. These RNAs, which compromise a substantial portion of the transcriptome, are packaged into granules and transported to subcellular compartments such as synapses, dendrites, and axons, and are critical for neuronal connectivity and communication. mcDETECT addresses this gap by identifying the spatial locations of RNA granules and reconstructing their transcriptome profiles from 3D subcellular mRNA distributions. In simulated data, mcDETECT demonstrated accurate RNA granule detection with minimal false positives. Using a public Xenium 5K dataset, we further showed that RNA granules have clear biological relevance: their presence is quantitatively linked to cellular compartments, and their spatial organization relative to neurons reveals regulatory layers that somatic expression alone cannot capture.

To demonstrate mcDETECT’s utility in disease studies, we designed a specialized gene panel and generated MERSCOPE data from WT and AD mouse brains. Analysis of this in-house dataset revealed disease-associated granule subtypes and brain region-specific distributional changes between WT and AD. These alterations also uncovered neuronal substates that are diminished or expanded in the AD model, highlighting disruption of the excitatory–inhibitory balance during early AD progression. Such findings are not detectable through somatic expression analysis alone, positioning mcDETECT as a valuable and complementary tool for exploring the hidden dark transcriptome.

A key advantage of mcDETECT is that it does not require extensive marker coverage; fewer than ten marker genes are sufficient to identify most granules. This flexibility enables mcDETECT to be applied to iST datasets with diverse gene panels, including those with limited granule marker coverage. To demonstrate this, we analyzed four WT mouse brain datasets generated by different platforms, including Xenium 5K, MERSCOPE, MERFISH, and CosMx, and consistently identified RNA granules with similar size, distribution, and gene enrichment patterns. This capability facilitates re-analysis of existing datasets and provides deeper insight into RNA granule changes across various neurodegenerative diseases.

Despite its utility, mcDETECT has several limitations that suggest directions for future improvement. First, our method identifies RNA granules based on RNA spatial distribution, without detecting the associated RBPs. It cannot distinguish between RNA granules and clusters of RNAs localized within the same distal compartments, both of which are biologically meaningful structures. Currently, there are no techniques capable of simultaneously identifying RBPs and RNA molecules in a spatial context. Further investigation will be needed as such technologies continue to evolve. A second challenge is the inherently sparse nature of RNA granule expression. As reported previously, each granule typically contains only a small number of transcript copies. Consequently, RNA granule profiles are much sparser than those derived from somatic expression, leading to weaker signals for certain genes and complicating the identification of robust subtype-specific patterns. Finally, because this work is a computational method-focused study, our proof-of-concept comparison of early-stage AD and WT mouse brains was limited to a single sample per group. Future studies with larger sample sizes will be essential to validate our findings and to fully leverage mcDETECT for identifying diverse RNA granule subtypes and neuronal substates across different stages of AD. We anticipate that mcDETECT will serve as a novel and valuable tool for advancing our understanding of the dark transcriptome. Ultimately, its findings may support early diagnosis and guide the development of RNA granule-targeted precision therapies for brain diseases.

## Acknowledgements

J.H. was supported by Emory AI Humanity Initiative, Georgia Clinical & Translational Science Alliance, and R35GM159880. B.Y. was supported by R01MH117122, R01AG062577, R01AG064786, R01AG078937, and R01NS118819.

## Author Contributions

This study was conceived and led by J.H. and B.Y. C.Y. designed the model and algorithm with input from J.H., implemented the mcDETECT software, and led data analyses with evaluation input from V.G.C., Hailing S., S.D., J.Y., P.J., and B.Y. K.P., Hongshun S., H.-L.V.W., F.W., and R.L. generated the MERSCOPE mouse brain data. Y.L. contributed to part of the data analysis. J.H. and C.Y. wrote the paper with feedback from all other co-authors.

## Methods

### Input data

mcDETECT is designed to accommodate a wide range of iST datasets generated by multiple iST platforms. Most iST platforms provide a file that records each transcript together with its gene identity and the (*x,y,z*) coordinates of the associated mRNA molecule, which serves as the primary input for identifying RNA granule locations. Some platforms also include a binary label indicating whether each mRNA molecule overlaps with a cell soma. In addition, mcDETECT requires a user-defined list of granule markers and negative control markers enriched in neuronal nuclei. We have summarized a set of distal RNA granule markers (**Supplementary Table 2**) and negative controls (**Supplementary Table 6**), and users may retain those that are present in their iST dataset.

#### Generating in-soma labels

If the input transcript file lacks binary in-soma labels for transcripts, mcDETECT performs an embedded pre-processing step to generate them based on DAPI images. First, image segmentation with adaptive thresholding is performed on the DAPI images to identify nuclear masks. Morphological dilation is then applied using a 3 × 3 square kernel for two iterations, which expands the nuclear masks outward to approximate cell somata. Next, the transcript coordinates are registered to image coordinates, allowing all mRNA transcripts to be overlaid onto the resulting binary DAPI image. From this registration, the relative position of each mRNA molecule with respect to the soma can be determined, enabling the assignment of binary in-soma labels.

### mcDETECT algorithm

mcDETECT is an end-to-end framework that processes transcript files from the iST platform along with user-defined granule markers, returning the spatial characteristics and transcriptome profiles of individual RNA granules. The complete mcDETECT algorithm consists of the following four steps:

#### Step 1: Density-based clustering on each single marker to identify extrasomatic aggregates

The fundamental step of mcDETECT is to identify extrasomatic aggregates of mRNAs in the input transcript file using a clustering strategy inspired by Density-Based Spatial Clustering with Applications of Noise^49^. It operates based on two predefined parameters: the search radius *r* and the minimum number of samples *m*. Denote *T* as the set of mRNAs from a granule marker, where each mRNA *t*_*i*_ ∈ *T* has 3D spatial coordinates (*x*_*i*_,*y*_*i*_,*z*_*i*_). We first calculate the Euclidean distance between all mRNA pairs *t*_*i*_, *t*_*i*_ *∈ T* by:

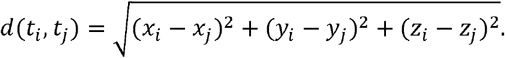

The neighbors of *t*_*i*_ are defined as the mRNAs falling within its neighborhood of radius *r*, and the number of its neighbors is recorded by:

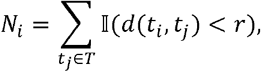

where II (·) is the indicator function. Next, mcDETECT classifies the mRNAs within the set *T* as core points, border points, and noise points according to the following criteria:

1. a point *t*_*i*_ is a core point if the number of its neighbors exceeds or equals the minimum number of samples *m*, i.e.,*N*_*i*_ ≥ *m*;
2. a point *t*_*j*_ is directly reachable from a core point *t*_*i*_ if it is within the neighborhood of *t*_*i*_, i.e., *d*(*t*_*i*_, *t*_*j*_) < *r;*
3. a point *t*_*j*_ is reachable from a core point *t*_*i*_ if there is a path { *t*_1_,*t*_2_, · · ·,*t*_*n*_} such that *t*_*i*_ = *t*_1_ and *t*_*j* =_ *t*_*n*_, where each point *t*_*i*+1_ is directly reachable from point *t*_*i*_ for *i* ∈ {1, · · ·,n −1};
4. a point *t*_*j*_ is a border point if the number of its neighbors is less than the minimum number of samples *m*, i.e., *N*_*j*_ *< m*, but it is reachable from a core point *t*_*i*_;
5. a point *t*_*k*_ is a noise point if it is not reachable from any other point.

A core point forms a cluster along with all points that are reachable from it, possibly consisting of other core points and border points, and each cluster represents a distinct aggregate of mRNAs. Based on this paradigm, mcDETECT effectively extracts the pattern where mRNAs aggregate while excluding the dissociated mRNAs.

The selection of the search radius *r* and the minimum number of samples *m* will affect the detection results. In this study, we set *r* to 1.5 μm, considering the typical size of RNA granules inferred from fluorescent puncta^29^. The minimum number of samples *m* is determined by mcDETECT through an automated procedure for each granule marker based on its transcript density. For a given granule marker, we first calculate its overall density *λ* as:

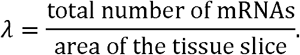

To assess whether the observed mRNA distribution deviates from randomness, we compare it against a Complete Spatial Randomness (CSR) process. Under CSR, the number of mRNAs within a circular region of radius *r* follows a Poisson distribution with mean *λπr*^2^. To distinguish true aggregates from random dispersion, we define the ideal minimum number of samples *m*^*\**^ as the value satisfying:

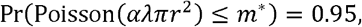

where *α* is a scaling factor used to adjust the density threshold for clustering. In this formulation, *m*^*^ represents the number of transcripts that would be expected only 5% of the time under CSR, making it a conservative threshold for identifying true aggregates. Due to the typically low values of *λ*in real datasets, we introduce *α* = 5 to enhance sensitivity, and we find that the results are not sensitive to further increases in *α*. Finally, in practice, the minimum number of samples *m* is determined as:

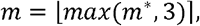

where we apply a flooring threshold of 3 to ensure that identified aggregates contain at least three mRNAs, and round *m* to its nearest integer. In conclusion, this data-driven parameter selection rocedure allows mcDETECT to automatically determine the best search radius *r* and minimum number of samples *m* to identify statistically significant aggregates for different input genes.

For each identified aggregate, mcDETECT constructs a minimum enclosing sphere that encompasses all associated mRNAs. The centroid and radius of this sphere serve as proxies for the location and size of the aggregate, respectively. Although the aggregates are initially defined using a single granule marker, the enclosing spheres allow for evaluating colocalization with other granule markers. The location, size, and granule marker composition are then recorded as spatial characteristics of the identified aggregates and are used for downstream filtering. Aggregates with an excessively large radius (4 μm) are discarded, as they likely represent cell fragments rather than distal RNA granules. In addition, mcDETECT evaluates t hein-soma ratio of each aggregate using the intracellular position of mRNAs. Specifically, let *B* = {*b*_1_,*b*_2_, · · ·,*b*_n_} be the set of all granule marker mRNAs within the enclosing sphere. An aggregate is discarded if:

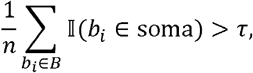

where *τ* is a user-defined threshold, whose value may vary depending on how in-soma labels are derived.

In this study, we set *τ* = 0.2 for iST datasets that provide in-soma labels per transcript. For datasets where in-soma labels are inferred from DAPI-based segmentation, we adopt a more conservative threshold of *τ* = 0.1 to account for potential inaccuracies.

#### Step 2: Integrating aggregates from multiple markers

To reveal the overall distribution of RNA granules informed by all granule markers, mcDETECT integrates aggregates across multiple markers while accounting for their spatial colocalization. The integration proceeds in the order of marker expression intensity, consistent with previous reports that highly expressed markers tend to colocalize with other genes^29^. This integration is performed using an iterative procedure based on the relative positions of spheres in 3D space. Specifically, let *s*_1_ and *s*_2_ represent two spheres derived from two different markers, with centroids *c*_1_ and *c*_2_, and radii *r*_1_ and *r*_2_, respectively. We first calculate the Euclidean distance between the centroids:

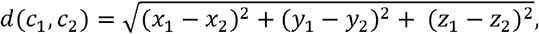

where (*x*_1_,*y*_1_,*z*_1_) and (*x*_2_,*y*_2_,*z*_2_) are the 3D spatial coordinates of *c*_1_and *c*_2_, respectively. Based on this inter-centroid distance, the relative positions of *s*_1_ and *s*_2_ are classified and handled as follows:

1. Disjoint: if *d* (*c*_1,_*c*_2_)≥ *r*_1_ + *r*_2,_the spheres do not overlap, and both *s*_1_ and *s*_2_ are retained.
2. Contained: if 0 ≤ *d* (*c*_1,_*c*_2_) ≥ | *r*_2_− *r*_1_|, the smaller sphere is considered contained within the larger one and is merged accordingly.
3. Intersecting: if | *r*_2_− *r*_1_| < *d* (*c*_1,_*c*_2_) < *r*_1_ *+ r*_2_, the spheres partially overlap. If the overlap is deemed significant, defined as *d* (*c*_1,_*c*_2_) < *ρ*·(*r*_1_ *+ r*_2_), where *ρ* is a threshold parameter, the mRNAs from *s*_1_ and *s*_2_ are merged, and a new enclosing sphere is computed. Otherwise, both aggregates are retained.

In this study, we set *ρ* = 0.2 to define “heavy overlap” and prevent merging of nearby but distinct RNA granules.

In mcDETECT, this integration procedure is applied iteratively across all pairs of granule marker-derived aggregates to ensure efficient consolidation of overlapping signals.

#### Step 3: Filtering based on negative control enrichment

Following multi-marker integration, mcDETECT performs an additional filtering step based on the enrichment of negative control markers. Based on previous studies, we select these marker genes to be enriched in neuronal nuclei compared to the cytoplasm^1,28,50^. For each aggregate, mcDETECT counts the number of negative control mRNAs within its associated minimum enclosing sphere. An aggregate is discarded if the ratio of negative control mRNAs to granule marker mRNAs exceeds 10%. Overall, this step helps reduce false positives caused by cell fragments or nonspecific mRNA accumulation. Aggregates that pass all screening steps, including size, in-soma ratio, and negative control enrichment, are retained as individual RNA granules.

#### Step 4: Constructing spatial transcriptome profiles for individual RNA granules

To build transcriptome profiles of the identified RNA granules, mcDETECT maps all mRNA transcripts from the iST dataset to the minimum enclosing spheres that represent individual RNA granules. For each granule, the number of mRNAs per gene is counted, resulting in a gene-by-granule count matrix. By combining this matrix with spatial coordinates, mcDETECT generates a comprehensive spatial transcriptome profile for RNA granules across the tissue section, which enables further investigations of granule heterogeneity and molecular alterations in neurodegenerative diseases.

### Downstream analyses

#### RNA granule subtyping

To classify the identified RNA granules into molecular subtypes, we use a predefined panel of granule-subtype-specific genes, including markers for pre-synaptic, post-synaptic, dendritic, and axonal compartments. We perform mini-batch K-Means clustering on the enrichment matrix of these genes across all granules, resulting in 15 initial granule clusters. By examining the enrichment patterns of the subtype-specific markers within each cluster, we annotate the clusters as representing granule populations enriched for pre-synaptic, post-synaptic, dendritic genes, or mixed signals.

#### Spatial domain assignment

To identify spatial domains in an iST dataset, we first generate a 2D grid over the tissue section, where each grid spot represents a 50 × 50 μm^2^ region. Next, we reconstruct spatial gene expression by mapping all mRNA transcripts onto these pseudo-spots. Using this spot-level gene expression matrix, we apply a spatial clustering method, SpaGCN^51^, to identify distinct spatial domains. These domains are subsequently annotated as major brain regions based on anatomic maps from the Allen Mouse Brain Atlas. Finally, the assigned brain region labels for each pseudo-spot are propagated to all cells and RNA granules within that spot, enabling region-specific spatial analysis at multiple biological scales.

#### Calculating RNA granule density

The 50 × 50 μm^2^ pseudo-spots are used as the smallest unit for summarizing granule density within the tissue region. The granule density at the spot level is calculated as the number of granules divided by the spot area. At the regional level, granule density is computed by averaging the spot-level densities across all spots within a given brain region. To evaluate the agreement between regional synaptic granule densities estimated by mcDETECT and the synapse density derived from the genetic labeling approach from age-matched mice, we use a weighted Spearman correlation coefficient^52,53^, accounting for the brain region area. Specifically, let *A* = {*a*_1_,…,*a*_*n*_} and *B* = {*b*_1_,…,*b*_*n*_}denote the granule densities across *n* brain regions obtained from the two approaches, and let {*w*_1_,…,*w*_*n*_} represent the corresponding areas of these brain regions. We begin by computing the ranks *R*_*A*_ = {*R*(*a*_1_),…, *R*(*a*_n_)} and *R*_*B*_ = {*R*(*b*_1_),…, *R*(*b*_n_)}. The weighted mean rank for group *A* is calculated as 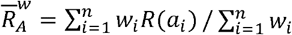, and similarly for 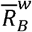 The weighted Spearman correlation is then calculated as:

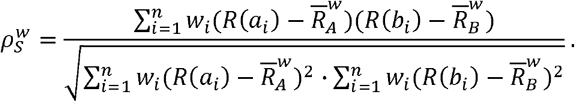

This weighted correlation provides a more accurate similarity measure between regional granule densities by assigning greater importance to brain regions with larger spatial coverage.

#### Granule context embedding and neuronal substate identification

To characterize the local microenvironment of each neuron, we compute a granule context embedding that captures the subtype composition of nearby RNA granules in a spatially weighted manner. For each neuron *i*, we first identify all its neighboring RNA granules within a predefined radius *r* = 10 μm. Let *N*_*i*_ denote the set of neighboring granules, and let each granule *j* ∈ *N*_*i*_ have subtype and *s*_*j*_ ∈ {1,…,*S*} spatial distance *d*_*ij*_ from neuron *i*. The contribution of each granule is modulated by an exponential decay kernel:

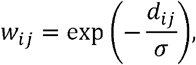

with the decay parameter *σ* fixed at a default of 5 μm in this study. Granule subtypes are one-hot coded into vectors ***z***_*j*_ ∈ {0,1}^*S*^, and a weighted sum across all neighboring granules is computed as:

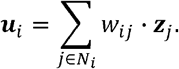

To obtain the final embedding, this vector is normalized to yield a probability distribution over granule subtypes:

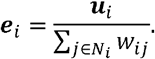

If a neuron lacks neighboring granules (i.e., *N*_*i*_ = Ø), it is assigned a one-hot embedding corresponding to a padding category (“others”) appended to the subtype list. This procedure yields a subtype composition vector ***e***_*i*_ *∈* ℝ ^*S*^ for each neuron, capturing the local granule-defined microenvironment.

To uncover neuronal substates with distinct granule contextual signatures, we cluster the granule context embeddings using Latent Dirichlet Allocation (LDA). Neurons within each brain region of interest are first subsetted, and their embeddings 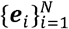 are modeled as samples from a *K*-component Dirichlet mixture. Each neuron is then assigned to a neuronal substate based on its maximum posteriori topic assignment:

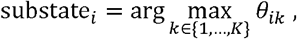

where *θ*_*ik*_ denotes the inferred mixture proportion of component *k*. The resulting cluster labels are encoded as ordered categorical variables. This combined embedding-clustering approach provides a principled framework for defining fine-grained neuronal substates informed by local RNA granule subtype composition.

### Simulation

#### Simulation of a single granule marker

The simulation is performed over a 2000 × 2000 μm^2^ area with a thickness 10.5 μm, mimicking the scale of the Xenium platform. To model the subcellular mRNA distribution of a single granule marker, we assume it consists of three spatial components: complete spatial randomness (CSR), intrasomatic aggregation, and extrasomatic aggregation. The simulation procedures for generating 3D coordinates (*x,y,z*) for each mRNA follow the methodology described by Diggle^54^, as detailed below.

1. CSR. The CSR pattern is modeled using a homogeneous Poisson process, where the *x,y*, and *z* coordinates are independently sampled from uniform distributions within the simulation bounds.
2. Intrasomatic aggregation. This aggregation pattern is modeled using a Poisson cluster process. First, cell soma centroids are generated via a homogeneous Poisson process. A random number of mRNAs are then assigned to the vicinity of each centroid, forming clusters around them. The number of mRNAs per soma is sampled from a Poisson distribution. Distances *d* between mRNAs and their corresponding soma centroids are independent and identically distributed (i.i.d.) exponential random variables. The directional angles of these distances are also i.i.d., determined by two uniform random variables: *φ*, the polar angle, ranging within (0,*π*), and *ϕ*, the azimuthal angle, ranging within (0,2*π*). Consequently, the3D offsets from soma centroids to mRNAs are given by:

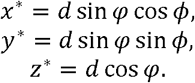

The final coordinates (*x,y,z*) of each mRNA are then calculated by adding the respective offsets to the soma centroid location. Repeating this for all somata yields the full clustering pattern within the simulated area. To simulate intrasomatic labeling, a binary in-soma label is assigned to each mRNA based on a Bernoulli distribution with success probability *p*, representing the likelihood that an mRNA resides within a soma. For each cluster, *p* is independently sampled from a Beta(8, 2) distribution, producing an overall mean in-soma ratio of 0.8. Additionally, the distances *d* are sampled with a mean of 3.5 μm, reflecting typical soma sizes.

Extrasomatic aggregation. This follows the same procedure as intrasomatic aggregation, with two key differences. First, the in-soma probability *p* is sampled from a Beta(2, 8) distribution, resulting in a lower average in-soma ratio of 0.2. Second, the distances *d* are sampled with a lower mean of 1 μm, reflecting reported RNA granule sizes.

In the final step, the CSR points, intrasomatic aggregates, and extrasomatic aggregates are overlaid within the simulated area to form the complete spatial pattern. A 2D version of the simulation can be generated by omitting the z-dimension.

#### Simulation of multiple granule markers

We consider three granule markers in the multi-marker scenario. As in the single-marker setting, each marker’s subcellular mRNA distribution follows one of three spatial patterns: CSR, intrasomatic aggregation, or extrasomatic aggregation, with the latter two distinguished by differences in in-soma ratios and aggregate sizes. Additionally, we account for the colocalization patterns of multiple granule markers within somata and distal neuronal compartments. The simulation procedures for this scenario are detailed below:

1. CSR. CSR patterns are simulated independently for each granule marker.
2. Intrasomatic aggregation. This aggregation pattern is modeled using a multi-sample Poisson cluster process. First, we generate cell soma centroid locations using a homogenous Poisson process. Then, we determine which of the three markers are expressed in each soma. Specifically, let *K* be a categorical random variable representing the number of markers expressed in a given soma, drawn from the following probability mass function (*pmf*):

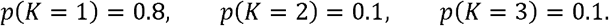

Given the value of *K*, we randomly select the corresponding markers and independently simulate each marker’s mRNA cluster following the procedures described in the single-marker setting. This results in clusters with diverse marker compositions.
3. Extrasomatic aggregation. This pattern follows the same procedures as intrasomatic aggregation, with adjustments to reflect different biological characteristics. In particular, the in-soma ratios are lower, aggregate size are smaller, and the *pmf* of *K* is modified to:

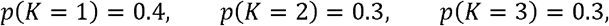

reflecting a higher degree of granule marker colocalization in distal neuronal compartments.

For both single-marker and multi-marker scenarios, we apply mcDETECT to the simulate mRNAs to recover the centroids of extrasomatic aggregates, which serve as ground truth. To evaluate the detection performance, we calculate the accuracy, precision, and recall based on whether these centroids are encompassed within the spheres identified by mcDETECT. The full simulation setup, including marker densities, the number of aggregates, and parameters for all distributions, is summarized in **Supplementary Table 4**.

### Regional connectivity map

Oh *et al*. ^55^reported a whole-brain connectivity matrix based on axonal projection strengths measured using enhanced green fluorescent protein (EGFP)-expressing adeno-associated viral vectors. From their connectivity matrix, we select the subset of brain regions available in our iST datasets, resulting in a square matrix where each element represents the connectivity strength from a source region (rows) to a target region (columns). We then calculate ex_total_ and im_total_, representing the total export and import strengths of each region, by summing the corresponding rows and columns. Based on these values, we defined an export-to-import score (E/I score) for each region as:

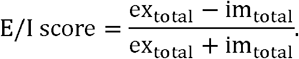

This score, ranging from −1 to 1, represents the relative role of a brain region in terms of sending or receiving axonal projections. An E/I score closer to 1 indicates that the region primarily functions as an outputting unit, whereas a score closer to −1 suggests it predominantly serves as an input-receiving unit.

### MERSCOPE data generation

#### Animals

5xFAD transgenic mouse (Jax Strain# 034848) and C57BL/6J (Jax Strain# 000664) were obtained from Jackson Laboratory, bred and maintained at Emory University School of Medicine. The whole brains were collected at 8 weeks-age. All animal procedures and protocols were approved by the Emory University Institutional Animal Care and Use Committee (Protocol ID: PROTO201800040).

#### Transcardinal Perfusion

Mice were anesthetized with isoflurane and perfused with 25–30 mL cold phosphate buffered saline (1× PBS) at a constant flow rate of 5 mL/min for about 6–8 min. After perfusion, the whole brain was collected, transferred onto pre-chilled mouse brain matrices on ice and spliced around the region of interest. Then the brain slice was transferred into a plastic cryo-mold filled with prechilled OCT compound on ice and immediately immersed into prechilled isopentane into liquid nitrogen for about 1– 1.5 min. After rapid freezing, placed the frozen tissue on dry ice, foil wrapped and stored at −80 °C freezer for long term.

#### Tissue preparation and hybridization for MERSCOPE

A 10 μm thick cryosection of a fresh frozen brain region from an 8-weeks age male 5xFAD heterozygous mouse and age-matched control wild-type littermates were prepared. The tissue section was mounted on the MERSCOPE slide stored at −20 °C for overnight, air dry for 5 min at RT and gently rinsed twice with 1× PBS for 5 min. Then, incubated for 15 min in 4% Pfa at RT and rinsed three times with 1× PBS for 5 min each. The section was stored in 70% ethanol at 4 °C.

Following the manufacturer’s protocol manual (Vizgen, *MERSCOPE Fresh and Fixed Frozen Tissue Sample Preparation User Guide*, 91600002, Rev F, 2023), the section was prepared for MERSCOPE probe hybridization. The MERSCOPE slide was removed from 70% ethanol, transferred into a small sterile petri dish with the tissue side facing up, and briefly rinsed once with 5 mL of Sample Prep Wash Buffer. Then, we aspirated the Sample Prep Wash Buffer and incubated the section in 5 mL of Formamide Wash Buffer at 37 °C for 30 min in a humidified incubator. After 30 min, the Formamide Wash Buffer was carefully aspirated from all around the section, and we added 50 μL of our predesigned 300-gene panel probe directly onto the tissue section. Then, we used tweezers to peel off the 2 × 2 cm^2^ parafilm to carefully cover and seal the probe on the tissue section. We then sealed the petri dish and incubated the section at 37 °C for 36–48 hours in a humidified incubator.

After probe hybridization, the section was incubated twice in 5 mL of Formamide Wash Buffer at 47 °C for 30 min each. While incubating, we prepared the Gel Embedding Solution containing Gel Embedding Premix, 10% w/v ammonium persulfate solution, and N,N,N’,N’,-tetramethylethylethylenediamine (5 mL per sample) and placed it on ice until ready to use. Then, we prepared the Gel Coverslip by applying 100 μL of Gel Slick Solution onto the Gel Coverslip for at least 10 min at room temperature. After the second Formamide Wash Buffer incubation, we briefly washed the section with 5 mL of Sample Prep Wash Buffer. From the 5 mL of freshly prepared Gel Embedding Solution, we retained 100 μL and added the remaining solution into the petri dish containing our section, incubating it for 1 min at room temperature. Then, we transferred the Gel Embedding Solution into a waste 50 mL tube to monitor gel formation and applied 50 μL of the retained Gel Embedding Solution directly onto the tissue section. We gently wiped the coverslip to remove any residue, then lifted the coverslip using tweezers, ensuring that the Gel Slick-treated side faced downward toward the tissue. Using another tweezer to guide and angle the coverslip, we slowly mounted it onto the tissue section without trapping air bubbles. We adjusted the position of the coverslip so that it was centered on the MERSCOPE slide. Excess Gel Embedding Solution was removed by gently tapping on the coverslip and carefully aspirating the excess solution from the sides. Without disturbing the petri dish, we allowed 1.5 hours of incubation at room temperature for gel formation. After that, we used the sharpened tip of a hobby blade to carefully lift the coverslip and proceeded to the next step of clearing for non-resistant fresh frozen tissue.

We used prewarmed Clearing Premix at 37 °C for 30 min and prepared a 5 mL Clearing Solution containing the Clearing Premix and 50 μL of Proteinase K. The Clearing Solution was then added to the petri dish containing the section, after which we sealed the petri dish and placed it in a humidified 37 °C cell culture incubator for 24 hours or until the tissue section became transparent.

#### MERSCOPE imaging

To prepare our tissue section for imaging, we first stained it with DAPI and PolyT stain. We began by warming a water bath to 37 °C, ensuring the water level was adequate but did not cover the valve of the Imaging Cartridge. The Imaging Cartridge was placed in the water bath for 60 min to allow it to thaw. Additionally, the DAPI and PolyT Staining reagent were prewarmed for 10 min in a 37 °C water bath. Once the tissue was cleared and became transparent, we aspirated the Clearing Solution and washed the section twice with 5 mL of Sample Prep Wash Buffer. Then, we added 3 mL of prewarmed DAPI and PolyT Staining reagent and allowed it to incubate for 15 min on a rocker. This was followed by a wash with 5 mL of Formamide Wash Buffer, with a 10 min incubation. The section was washed again with 5 mL of Sample Prep Wash Buffer and left in the buffer until the MERSCOPE instrument was ready for imaging.

For MERSCOPE instrument operation and data acquisition, we followed the manufacturer’s protocol manual (Vizgen, *MERSCOPE Instrument User Guide*, 91600001, Rev H, 2023). Before starting the MERSCOPE run, we first washed the instrument according to the manual’s instructions. In this experiment, we did not perform the MERSCOPE Cell Boundary Stain or any additional antibody staining. We ensured that the Imaging Cartridge was completely thawed, then slowly inverted the Cartridge ten times to mix the reagents inside. The Cartridge was activated by piercing the foil at the Cartridge Activation Port with a clean pipette tip. Using a 1 mL pipette set to 1 mL, we carefully added all of the Imaging Activation Mix into the bottom of the Cartridge via the Activation Port and plunged the pipette ten times at a moderate speed to thoroughly mix the solution inside. Then, using a serological pipette, we carefully layered 15 mL of mineral oil over the liquid inside the Imaging Cartridge. The Imaging Cartridge was inserted into the MERSCOPE instrument, with the barcode side facing the front of the instrument.

Next, we assembled the MERSCOPE Flow Chamber parts along with the MERSCOPE Slide containing the tissue section in the correct orientation. The Flow Chamber was locked in place, and image acquisition proceeded according to the manual’s instructions. The total MERSCOPE imaging time for this experiment was approximately 24–36 hours, followed by data acquisition.

## Data availability

We analyzed the following publicly available iST datasets: (1) Xenium 5K mouse brain data (https://www.10xgenomics.com/datasets/xenium-prime-fresh-frozen-mouse-brain); (2) CosMx mouse brain data (https://nanostring.com/products/cosmx-spatial-molecular-imager/ffpe-dataset/cosmx-smi-mouse-brain-ffpe-dataset/); (3) MERFISH mouse brain data (https://info.vizgen.com/mouse-brain-map). We also analyzed two in-house MERSCOPE mouse brain datasets, which have been uploaded to GEO (accession number: GSE292210) with restricted access. They will be made publicly available upon acceptance of the manuscript. Details of the datasets analyzed in this paper are described in **Supplementary Table 1**.

## Software availability

An open-source implementation of the mcDETECT algorithm can be downloaded from GitHub: https://github.com/chen-yang-yuan/mcDETECT.

