## Supplementary Information for "Uncovering the dark transcriptome in polarized neuronal compartments with mcDETECT"

**with mcDETECT**

Chenyang Yuan, Krupa Patel, Hongshun Shi, Hsiao-Lin V. Wang, Feng Wang, Ronghua Li, Yangping Li, Victor G. Corces, Hailing Shi, Sulagna Das, Jindan Yu, Peng Jin, Bing Yao^*^ & Jian Hu^*^

**Table of Contents**

**Supplementary Table 1.** Datasets analyzed in this manuscript.

**Supplementary Table 2.** Marker genes used for RNA granule detection and subtyping.

**Supplementary Table 3.** Marker genes used for RNA granule detection in each dataset.

**Supplementary Table 4.** Simulation settings.

**Supplementary Table 5.** Customized gene panel in the MERSCOPE experiment (separate file).

**Supplementary Table 6.** List of negative control markers enriched in neuronal somata (separate file).

**Supplementary Note 1.** Simulation study in the 2D setting.

**Supplementary Note 2.** Enriched genes and pathways in cortical neuronal substates 1 & 2 in Xenium 5K.

**Supplementary Note 3.** Enriched genes and pathways in other cortical neuronal substates in Xenium 5K.

**Supplementary Note 4.** RNA granule count discrepancy between Xenium 5K and MERSCOPE.

**Supplementary Note 5.** Synaptic granule density in the MERSCOPE WT sample.

**Supplementary Note 6.** Disease-associated alteration of other granule subtypes in MERSCOPE.

**Supplementary Note 7.** Enriched genes and pathways in other cortical neuronal substates in MERSCOPE.

**Supplementary Note 8.** Enriched genes and pathways in hippocampal neuronal substates in MERSCOPE.

**Supplementary Note 9.** Removing top-expressed markers in RNA granule detection in Xenium 5K.

**Supplementary Table 1.** Datasets analyzed in this manuscript.

| **Tissue** | **Data Source** | **Protocol** | **Mouse Model** | **Dataset Dimensions** | **Transcripts Count** |
| --- | --- | --- | --- | --- | --- |
| Mouse brain hemisphere (coronal) | 10x Genomics (https://www.10x  genomics.com/  datasets/xenium-prime-fresh-frozen-mouse-brain) | Xenium 5K, fresh frozen | C57BL/6, male, 9 weeks | 5006 genes, 63173 cells, 14892 spots* | 186743974 transcripts, 70468619 (37.74%) unmapped to cells |
| Mouse brain hemisphere (coronal) | In-house | MERSCOPE, fresh frozen | C57BL/6J, female, 8 weeks | 290 genes, 104482 cells, 17667 spots | 103398068 transcripts, 59168101 (57.22%) unmapped to cells |
|  |  |  | 5xFAD, male, 8 weeks | 290 genes, 75848 cells, 12604 spots | 68876647 transcripts, 42839701 (62.20%) unmapped to cells |
| Mouse brain hemisphere (coronal) | nanoString ( https://nanostring.com/products/cosmx-spatial-molecular-imager/ffpe-dataset/cosmx-smi-mouse-brain-ffpe-dataset/) | CoxMx, FFPE | C57BL/6, male, 7–8 weeks | 950 genes, 48180 cells, 16416 spots | 116084110 transcripts, 40588750 (34.96%) unmapped to cells |
| Mouse brain hemisphere (coronal) | Vizgen (https://info.vizgen.com/mouse-brain-map) | MERFISH, – | C57BL/6J, female, 6–8 weeks | 483 genes, 42089 cells, 15088 spots | 24045904 transcripts, – |

* For all datasets, the pseudo-spots are 50 × 50 μm^2^ square grids spanning the tissue slice (Methods).

**Supplementary Table 2.** Marker genes used for RNA granule detection and subtyping.

| **Subtype** | **Gene** | **Source Studies** |
| --- | --- | --- |
| Presynaptic (13) | *Bsn* | Cajigas *et al*.^1^, Hafner *et al.*^2^, Tarannum *et al.*^3^, Niu *et al*.^4^ |
|  | *Cplx2* | Cajigas *et al*., Niu *et al*. |
|  | *Gap43* | Cajigas *et al*., Niu *et al*. |
|  | *Nrxn1* | Cajigas *et al*., Niu *et al*. |
|  | *Slc17a6* | Niu *et al*., Reimer^5^ |
|  | *Slc17a7* | Cajigas *et al*., Hafner *et al.*, Niu *et al*., Zeng *et al*.^6^ |
|  | *Slc32a1* | Niu *et al*., Chaudhry *et al*.^7^ |
|  | *Snap25* | Cajigas *et al*., Niu *et al*. |
|  | *Stx1a* | Cajigas *et al*., Niu *et al*. |
|  | *Syn1* | Cajigas *et al*., Niu *et al*., Zeng *et al*. |
|  | *Syp* | Cajigas *et al*., Niu *et al*. |
|  | *Syt1* | Cajigas *et al*., Niu *et al*. |
|  | *Vamp2* | Cajigas *et al*., Niu *et al*. |
| Postsynaptic (13) | *Camk2a* | Cajigas *et al*., Tarannum *et al.*, Niu *et al*., Glock *et al*.^8^ |
|  | *Dlg3* | Cajigas *et al*., Niu *et al*. |
|  | *Dlg4** | Cajigas *et al*., Tarannum *et al.*, Niu *et al*., Zeng *et al*., Glock *et al*. |
|  | *Gphn* | Vlachos *et al*.^9^, Tretter *et al*.^10^ |
|  | *Gria1* | Cajigas *et al*., Zeng *et al*. |
|  | *Gria2* | Cajigas *et al*., Diering and Huganir^11^ |
|  | *Homer1* | Cajigas *et al*., Yoon *et al*.^12^ |
|  | *Homer2* | Cajigas *et al*., Niu *et al*., Zeng *et al*., Glock *et al*. |
|  | *Nlgn1* | Cajigas *et al*., Niu *et al*. |
|  | *Nlgn2* | Cajigas *et al*., Niu *et al*. |
|  | *Nlgn3* | Uchigashima *et al.*^13^ |
|  | *Shank1* | Cajigas *et al*., Niu *et al*., Zeng *et al*., Glock *et al*. |
|  | *Shank3* | Cajigas *et al*., Niu *et al*., Zeng *et al*., Glock *et al*. |
| Axonal (6) | *Ank3* | Niu *et al*., Sobotzik *et al.*^14^ |
|  | *Mapt* | Niu *et al*., Forrest *et al.*^15^ |
|  | *Nav1* | Sánchez-Huertas *et al.*^16^ |
|  | *Nfasc* | Zonta *et al.*^17^ |
|  | *Sptbn4* | Wang *et al.*^18^ |
|  | *Tubb3* | Latremoliere *et al.*^19^ |
| Dendritic (5) | *Actb* | Niu *et al*., Klein *et al.*^20^ |
|  | *Cyfip2* | Tarannum *et al.*, Niu *et al*. |
|  | *Ddn* | Tarannum *et al*, Niu *et al*. |
|  | *Map1a* | Niu *et al*., Takei *et al.*^21^ |
|  | *Map2* | Niu *et al*., DeGiosio *et al.*^22^ |

*: *Dlg4* is also considered a dendritic marker in this study.

**Supplementary Table 3.** Marker genes used for RNA granule detection in each dataset. Genes are ordered by their marginal contribution to the identified granules.

| **Dataset** | **Marker Genes** |
| --- | --- |
| Xenium 5K (24) | *Snap25, Camk2a, Slc17a7, Cyfip2, Map2, Syp, Syn1, Slc32a1, Vamp2, Mapt, Gria2, Gap43, Tubb3, Dlg4, Gria1, Bsn, Homer1, Slc17a6, Stx1a, Dlg3, Gphn, Shank3, Nhgn2, Nlgn3* |
| MERSCOPE (34) | *Camk2a, Cplx2, Slc17a7, Ddn, Syp, Map1a, Shank1, Syn1, Gria1, Gria2, Cyfip2, Vamp2, Bsn, Slc32a1, Nfasc, Syt1, Tubb3, Nav1, Shank3, Mapt, Stx1a, Nlgn2, Ank3, Nlhn1, Gphn, Nlgn3, Slc17a6, Gap43, Dlg3, Dlg4, Map2, Homer2, Homer1, Nrxn1* |
| CosMx (15) | *Snap25, Camk2a, Slc17a7, Syp, Syt1, Gap43, Cyfip2, Dlg4, Gria2, Slc32a1, Mapt, Homer1, Gria1, Nfasc, Gphn* |
| MERFISH (8) | *Slc17a7, Syt4, Adora1, Cnr1, Slc17a6, C1qb, Slc32a1, Slc17a8* |

– Blue: Not present in our compiled granule marker list (see **Supplementary Table 2**), but still related to synaptic, axonal, or dendritic function.

– Gray: Present in our compiled granule marker list, but not used for granule detection in this study due to limited marginal contribution (collectively accounting for less than 1% of all granules).

**Supplementary Table 4.** Simulation settings.

| **Overall Setting** | | | | | | | | | | | | | | |
| --- | --- | --- | --- | --- | --- | --- | --- | --- | --- | --- | --- | --- | --- | --- |
| Length (μm) | | Width (μm) | | Thickness(μm) | | | CSR points ratio | | | ESA points ratio | | | ISA points ratio | |
| 2000 | | 2000 | | 10.5 | | | 0.5 | | | 0.25 | | | 0.25 | |
| **Single-Marker Aggregate** | | | | | | | | | | | | | | |
| Marker type | Density (count/μm^2^) | | Count | | | Mean radius D (μm) | | | Beta distribution parameters for  in-soma label | | | | | |
|  |  |  | ESA | | ISA | ESA | | ISA | ESA | | | ISA | | |
| A | 0.08 | | 5000 | | 2000 | 1 | | 3.5 | (2, 8) | | | (8, 2) | | |
| B | 0.04 | | 3000 | | 1200 |  |  |  |  |  |  |  |  |  |
| C | 0.02 | | 2000 | | 800 |  |  |  |  |  |  |  |  |  |
| **Multi-Marker Aggregate** | | | | | | | | | | | | | | |
| Marker type | Density (count/μm^2^) | | Count | | | Mean radius D (μm) | | | Beta distribution parameters for  in-soma label | | | Probability of  {1, 2, 3} components | | |
|  |  |  | ESA | | ISA | ESA | | ISA | ESA | | ISA | ESA | | ISA |
| A | 0.08 | | 5000 | | 1000 | 1 | | 3.5 | (2, 8) | | (8, 2) | 0.4  0.3  0.3 | | 0.8  0.1  0.1 |
| B | 0.04 | |  |  |  |  |  |  |  |  |  |  |  |  |
| C | 0.02 | |  |  |  |  |  |  |  |  |  |  |  |  |

Abbreviations:

– CSR: complete spatial randomness

– ESA: extrasomatic aggregates

– ISA: intrasomatic aggregates

**Supplementary Table 5.** Customized gene panel in the MERSCOPE experiment (separate file).

**Supplementary Table 6.** List of negative control markers enriched in neuronal somata (separate file).

**Supplementary Note 1.** Simulation study in the 2D setting.

Contrasting the 3D setting, we conducted a similar simulation for each individual granule marker but suppressed the z-axis, treating all mRNAs as distributed in a 2D plane, as shown in **Fig. S1a**. The resulting detection accuracy, precision, and recall are displayed in **Fig. S1b**. Notably, across all three granule markers, the accuracy and precision remarkably drop compared to the 3D setting, since the randomly spread mRNA points became denser in 2D and mcDETECT falsely identified them as aggregates. Similarly, for the multi-marker scenario in 2D illustrated in **Fig. S1c**, mcDETECT generated higher false positives compared to 3D, resulting in reduced mean accuracy (93.2%) and precision (93.1%), as shown in **Fig. S1d**. In summary, the simulation study demonstrates that mcDETECT is more effective in 3D than in 2D at accurately recovering true RNA granule locations and excluding nuclear aggregates and random debris.

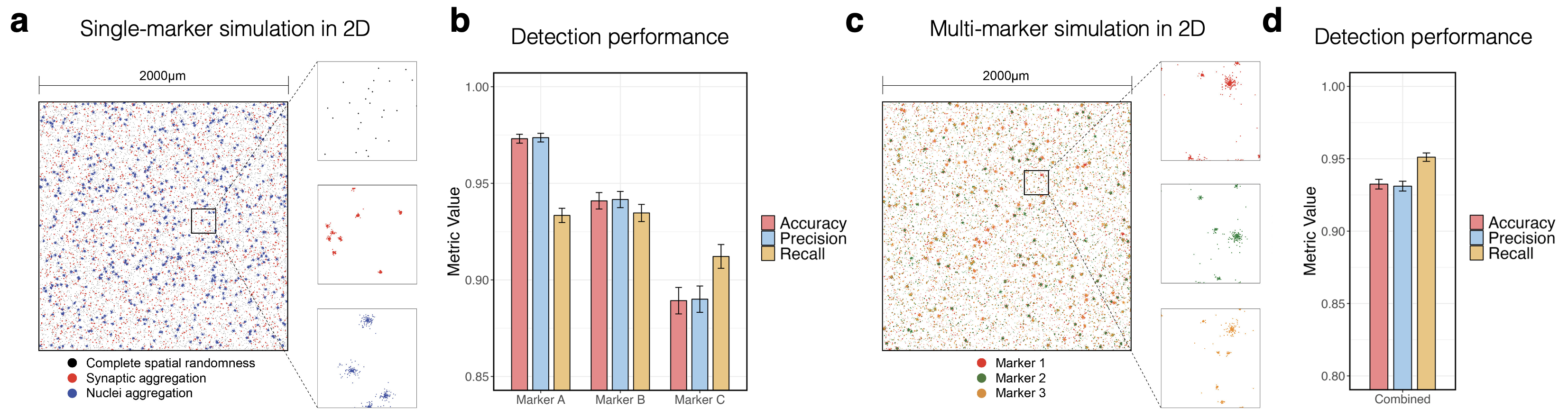

**Fig. S1.** **a.** Left: Simulation of a single granule marker in 2D space, where each dot represents a subcellular mRNA molecule. Black, red, and blue dots indicate complete spatial randomness (CSR), aggregate within distal compartments, and aggregate within somata, respectively. Right: Zoomed-in view of the three distribution patterns in a small region. **b.** Bar plot showing the detection accuracy, precision, and recall achieved by mcDETECT on the simulated 2D single-marker data across 200 simulation runs. Bar heights represent mean values, and error bars denote mean $\pm$ one standard deviation. **c.** Left: Simulation of three granule markers in 2D space, where each dot represents a subcellular mRNA molecule. Red, green, and orange dots indicate marker types A, B, and C, respectively. Right: Zoomed-in view of a small region stratified by marker type. **d.** Bar plot showing the detection accuracy, precision, and recall achieved by mcDETECT on the simulated 2D multi-marker data across 200 simulation runs. Bar plot definitions are the same as in **b**.

**Supplementary Note 2.** Enriched genes and pathways in cortical neuronal substates 1 & 2 in Xenium 5K.

A full interpretation of the granule context and the functional roles of Substates 1 and 2 is provided in the main text. Their respective enriched biological processes are visualized in **Fig. S2**.

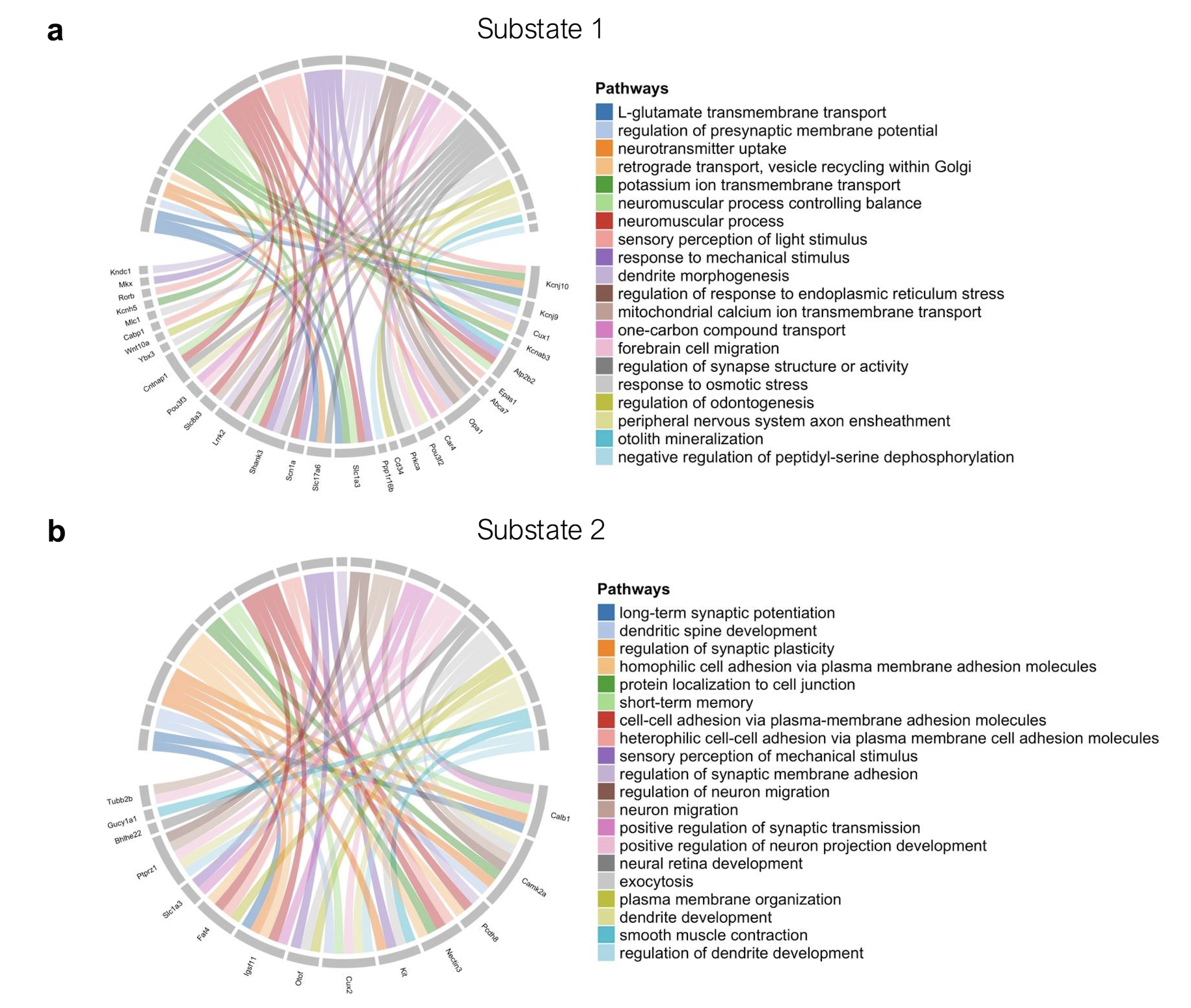

**Fig. S2.** Chord diagrams illustrating enriched biological processes and their associated genes in two representative neuronal substates: (a) Substate 1 and (b) Substate 2. Links represent gene–pathway associations based on GO enrichment analysis.

**Supplementary Note 3.** Enriched genes and pathways in other cortical neuronal substates in Xenium 5K.

In addition to Substates 1 and 2, we verified the association between granule context and the functional roles of other cortical neuronal substates in the Xenium 5K dataset (Substates 3–5). Specifically, we performed differential expression (DE) and Gene Ontology (GO) analyses based on their somatic expression, and the enriched genes and biological processes of each neuronal substate are summarized and visualized in **Fig. S3**.

Substate 3 neurons were enriched for surrounding pre-synaptic granules. This substate most likely represents a specialized subpopulation engaged in neurotransmitter release and vesicle dynamics. As shown in **Fig. S3a**, enrichment of biological processes such as synaptic vesicle maturation, priming, clustering, and exocytosis, along with regulation of GABAergic and glutamatergic transmission, suggests that these neurons are optimized for efficient presynaptic output as well as modulation of synaptic strength.

Substate 4 neurons were enriched for surrounding mixed-signal granules. This substate likely represents a hybrid subpopulation integrating presynaptic release mechanisms and postsynaptic plasticity responses. As shown in **Fig. S3b**, upregulation of gene programs involved in synaptic vesicle cycle, calcium-regulated exocytosis, vesicle docking, and long-term synaptic potentiation suggests that these neurons are engaged in both rapid neurotransmitter release and activity-dependent synaptic strengthening. Processes such as axo-dendritic transport and regulation of protein polymerization further indicate a role in structural remodeling and maintenance of neuronal homeostasis.

Substate 5 neurons exhibited enrichment of surrounding dendritic granules. This substate most likely represents a population specialized in postsynaptic integration and plasticity. As shown in **Fig. S3c**, upregulation of pathways such as dendritic spine development, synapse assembly, long-term synaptic potentiation, and regulation of synapse organization indicates that these neurons actively strengthen synaptic connections and shape memory-related processes. Additional processes involving calcium-regulated exocytosis and glutamatergic transmission further highlight their role in excitatory postsynaptic signaling and adaptive circuit remodeling.

Taken together, these functional profiles are consistent with the granule enrichment patterns of all cortical neuronal substates, demonstrating that granule-defined substates identified by mcDETECT represent functionally distinct neuronal populations, which are different from and complementary to those defined by somatic transcriptomes.

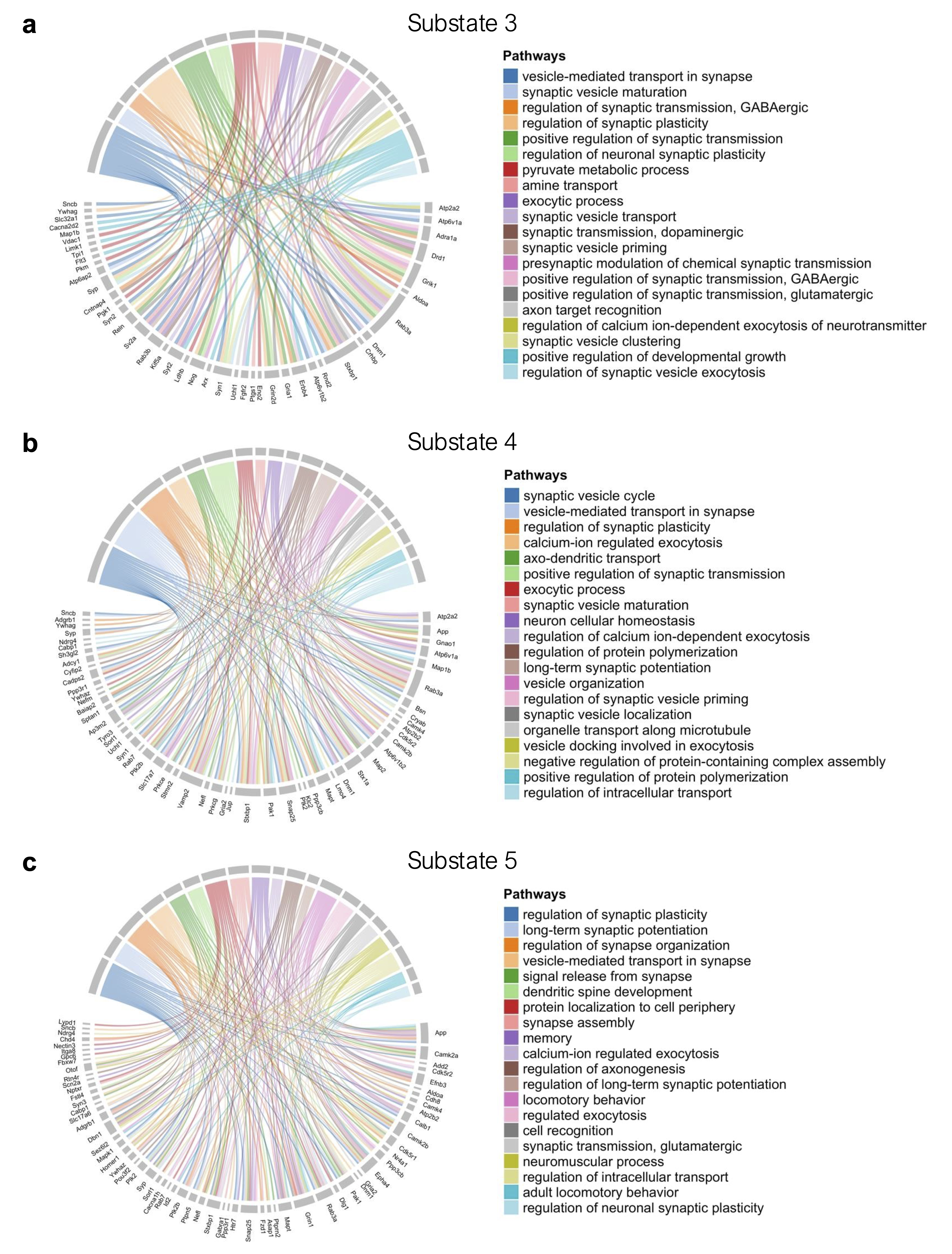

**Fig. S3.** Chord diagrams illustrating enriched biological processes and their associated genes in **(a)** Substate 3, **(b)** Substate 4, and **(c)** Substate 5 neurons. Links represent gene–pathway associations based on GO enrichment analysis.

**Supplementary Note 4.** RNA granule count discrepancy between Xenium 5K and MERSCOPE.

mcDETECT identified 170,501 RNA granules in the Xenium 5K dataset. In comparison, it identified 965,847 and 532,857 RNA granules in the MERSCOPE WT and AD samples, respectively. This discrepancy can be attributed to the following factors:

1. The input granule marker lists were different.
2. The MERSCOPE slices were positioned closer to the caudal end of the mouse brain than the Xenium 5K sample.
3. MERSCOPE generally exhibits higher mRNA capture efficiency than Xenium 5K. To quantify this difference, we first conducted a global comparison of all genes in both datasets. The average mRNA count per cell was 0.33 in Xenium 5K and 1.19 in the MERSCOPE WT sample, highlighting substantial platform variation. Next, we focused on the 166 genes shared between the two datasets and assessed their average mRNA count per spot to eliminate potential bias from cell segmentation. As shown in the overlaid bar plot in **Fig. S4**, most genes exhibited higher average mRNA counts per spot in MERSCOPE than in Xenium 5K, with an average fold increase of 3.27. For *Camk2a*, one of the strongest synaptic markers, the fold increase reached 3.89. Since mcDETECT relies on subcellular mRNA enrichment for granule identification, higher mRNA capture efficiency naturally results in a higher RNA granule count.

To account for differences in capture efficiency across experiments and platforms, we selected one dataset as an anchor. For each dataset being compared to the anchor, we first retained only the genes shared between the two datasets. We then calculated the mean transcript count per spot for both the anchor and the compared dataset and derived a scaling factor as the ratio of these two means (anchor / compared). When comparing RNA granule numbers, we multiplied the granule density from the compared dataset by this scaling factor. This adjustment enables more direct comparisons of granule numbers across datasets.

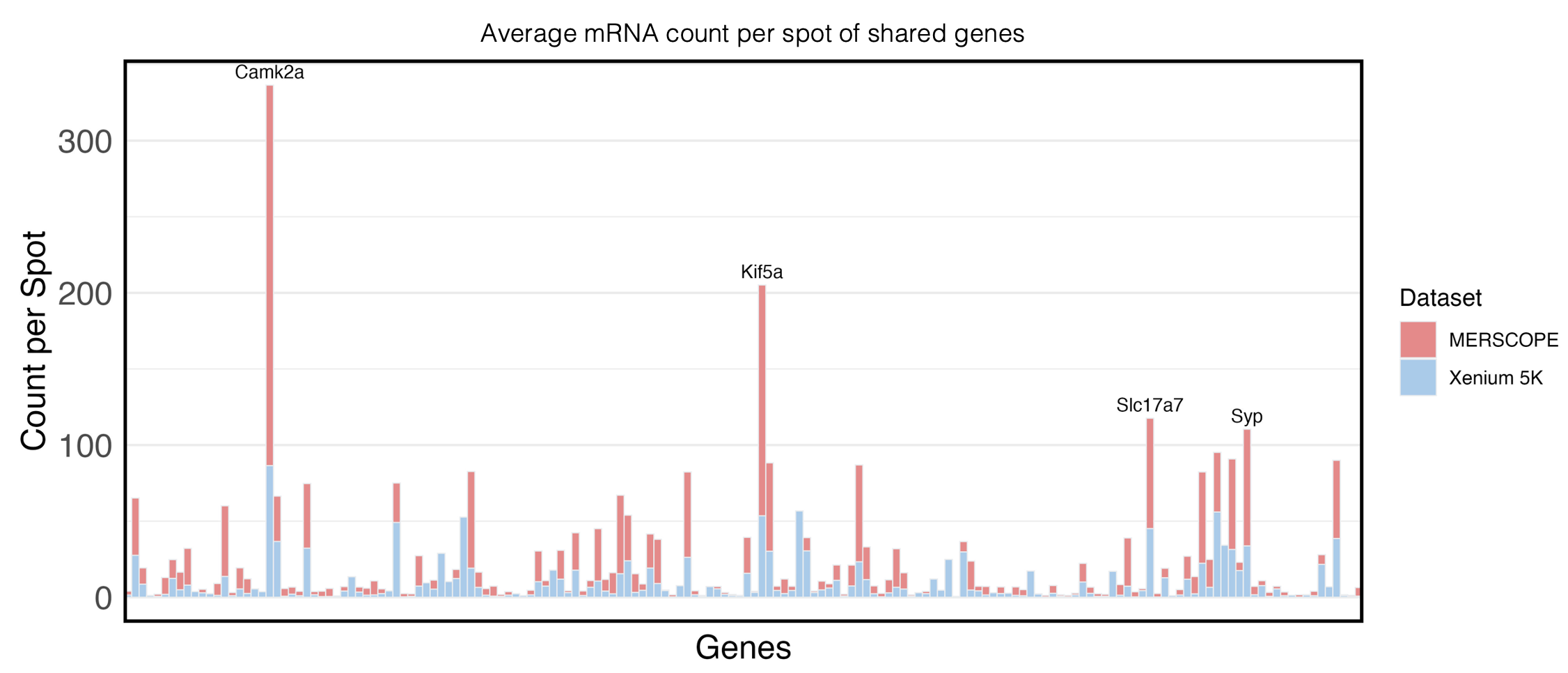

**Fig. S4.** Bar plot showing the average mRNA count per spot for the 166 shared genes between the Xenium 5K (blue) and the MERSCOPE WT sample (red). The bars are overlaid for direct comparison. Genes with an average mRNA count per spot exceeding 100 in the MERSCOPE WT sample are annotated.

**Supplementary Note 5.** Synaptic granule density in the MERSCOPE WT sample.

To demonstrate the biological relevance of the granule subtypes identified in the MERSCOPE WT sample, we quantified the density of synapse-associated granules (enriched for both pre- and post-synaptic genes) within each 50 × 50 μm² spot across nine major brain regions (**Fig. S5, left**). We compared these granule densities with synapse densities from an age-matched, similarly positioned WT mouse brain section, measured using volume electron microscopy and genetic labeling (**Fig. S5, right**), and found a weighted Spearman correlation coefficient of 0.96.

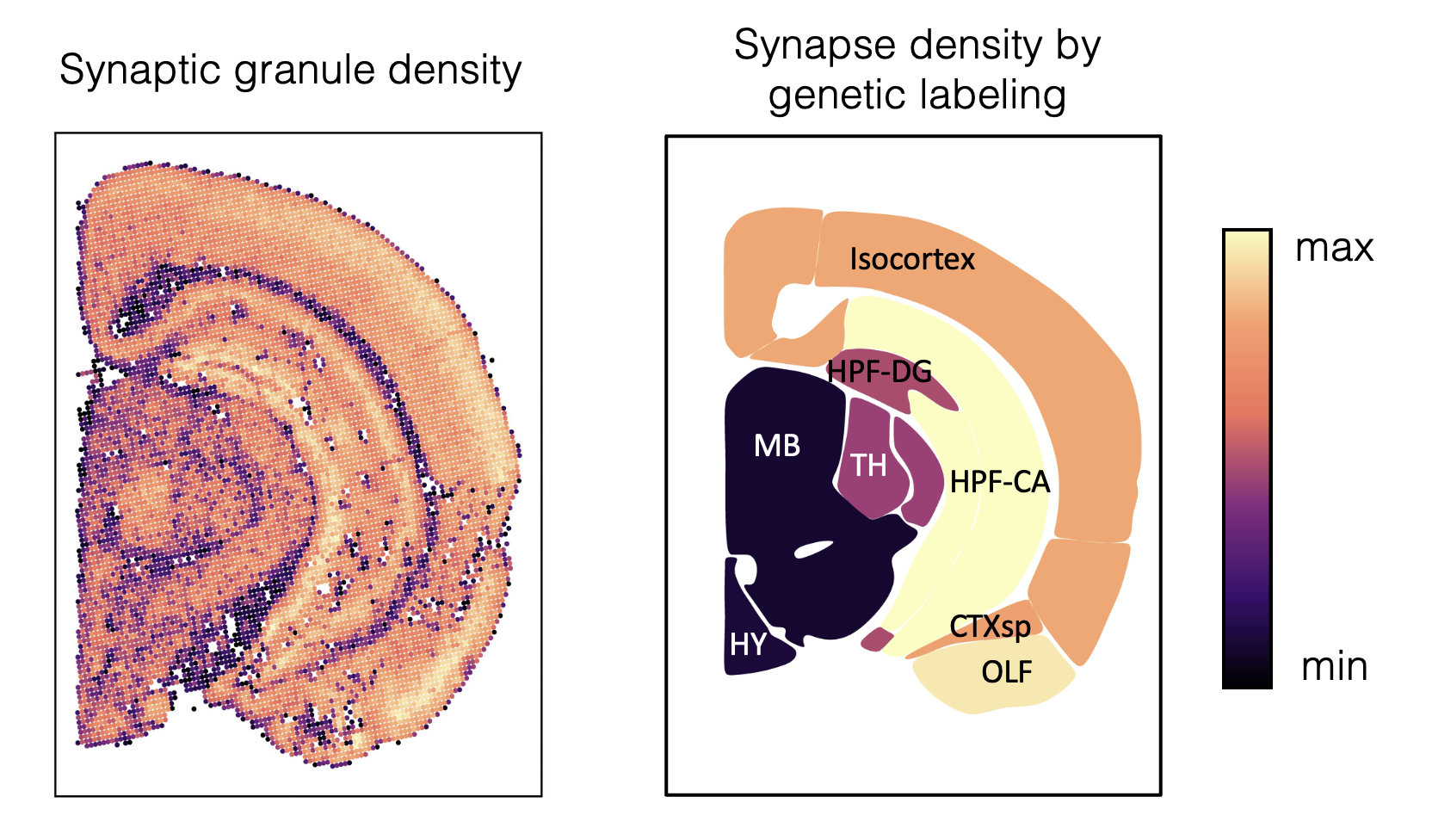

**Fig. S5.** Left: scatter plot showing the number of synaptic granules (i.e., pre- or post-synaptic) identified by mcDETECT in 50 × 50 μm^2^ spots spanning the tissue slice. Right: regional synapse densities estimated using volume electron microscopy and genetic labeling on a coronal slice corresponding to the approximate bregma position in **Fig. 4a**.

**Supplementary Note 6.** Disease-associated alteration of other granule subtypes in MERSCOPE.

In addition to granules enriched for pre- and post-synaptic genes, other subtypes, including those enriched for dendritic genes and those with mixed signals, also showed region-specific density changes in AD. As illustrated in **Fig. S6 (left)**, dendritic granules exhibited a marked decline in HPF-CA and TH, but a significant expansion in HPF-DG, HPF-SR, and MB. Mixed-signal granules, in contrast, displayed significant reductions across most brain regions, with the exception of HPF-DG and HPF-SR, where densities were significantly elevated (**Fig. S6, right**). However, given the non-uniform changes observed for dendritic granules and the ambiguous identity of mixed-signal granules, the biological significance of these alterations in AD remains unclear and warrants further investigation.

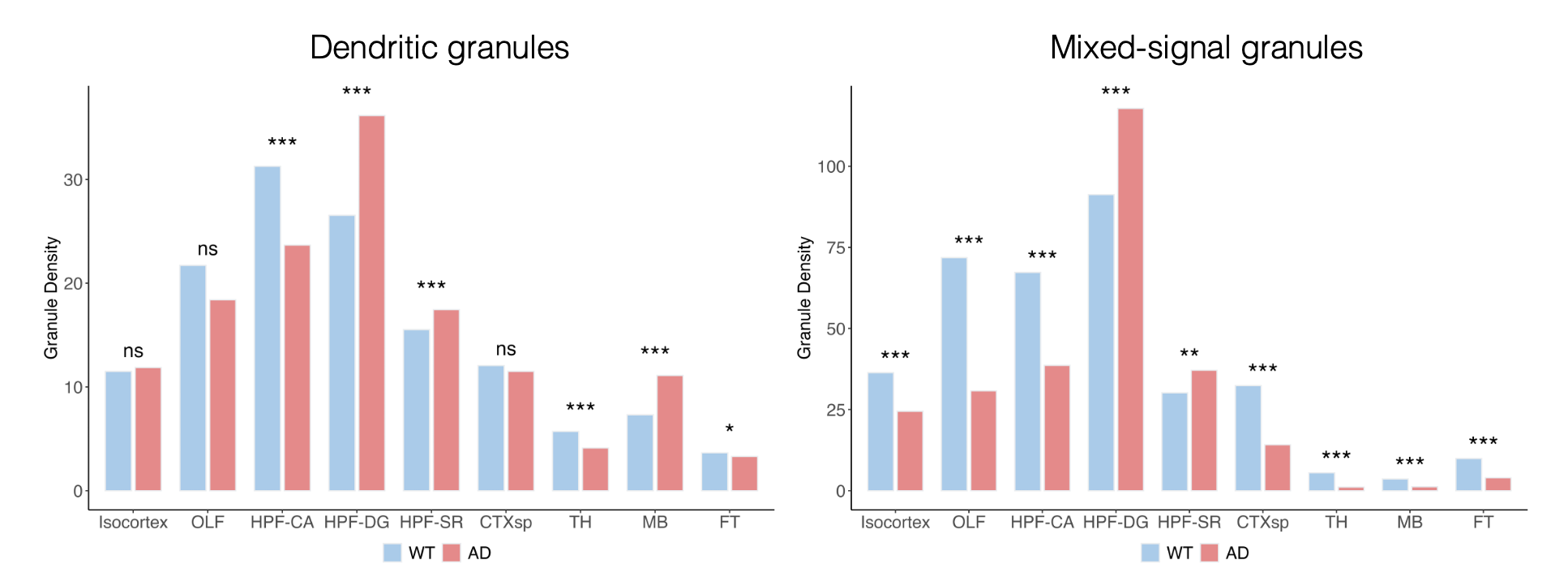

**Fig. S6.** Grouped bar plots comparing regional densities of dendritic (left) and mixed-signal granules (right) between WT and AD samples, after adjusting for differences in capture efficiency. Two-sample t-tests were performed on log-transformed spot-level granule counts within each brain region. ns: not significant; *: p < 0.05; **: p < 0.01; ***: p < 0.001.

**Supplementary Note 7.** Enriched genes and pathways in other cortical neuronal substates in MERSCOPE.

In addition to Substates 1 and 2, we evaluated the disease-associated alterations of other cortical neuronal substates (Substates 3–5) in the MERSCOPE dataset, as well as their biological relevance. Alterations in these neuronal substates revealed additional signatures of AD-induced circuit dysfunction.

Substate 3 neurons showed a significant increase in AD (**Fig. S7a, left**; two-sample t-test, p < 2.2 × 10⁻¹⁶) and were enriched for surrounding dendritic and ambiguously classified granules. This substate most likely represents a heterogeneous or less synapse-specialized population, supported by upregulated pathways such as neural crest cell migration, keratinocyte differentiation, and peripheral nervous system development (**Fig. S7a, right**). Instead, they may be characterized by stress-related, developmental, or non-canonical programs rather than by classical pre- or post-synaptic signaling functions.

Substate 4 neurons did not show disease-associated alterations (**Fig. S7b, left**; two-sample t-test, p = 0.097) and were enriched for surrounding post-synaptic granules. This substate exhibited upregulation of gene programs involved in the regulation of RNA biosynthetic pathways, transcription factor activity, calcium ion transport, and endoplasmic reticulum stress responses (**Fig. S7b, right**), suggesting that these neurons likely represent a population tuned toward postsynaptic responsiveness and activity-dependent regulation, engaged in maintaining protein homeostasis under high activity demands.

Substate 5 neurons showed a significant decrease in AD (**Fig. S7c, left**; two-sample t-test, p < 2.2 × 10⁻¹⁶) and were enriched for surrounding pre-synaptic granules. This substate most likely represents a population specialized in excitatory neurotransmission and presynaptic output. As shown in **Fig. S7c (right)**, enrichment of pathways such as ionotropic glutamate receptor signaling, synaptic vesicle exocytosis, SNARE complex assembly, and regulation of trans-synaptic signaling highlights their central role in fast synaptic transmission and synaptic plasticity. Additional pathways, including regulation of dendrite morphogenesis and small GTPase-mediated signaling, suggest structural adaptability and dynamic control over neurotransmitter release.

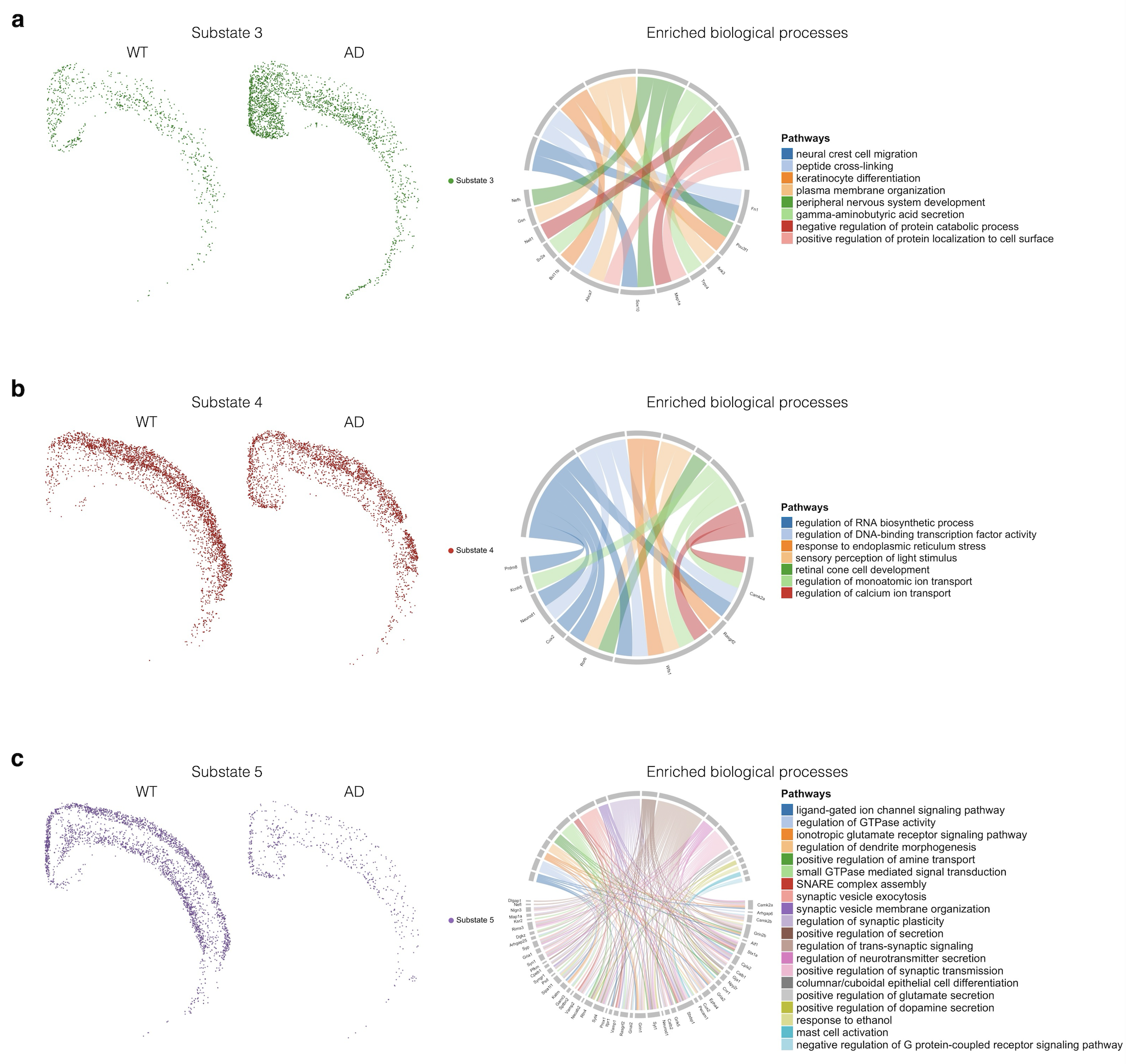

**Fig. S7.** **a.** Left: Scatter plot showing the distribution of Substate 3 neurons in WT and AD samples. Right: Chord diagram illustrating enriched biological processes and their associated genes in Substate 3 neurons. Links represent gene–pathway associations based on GO enrichment analysis. **b.** Left: Scatter plot showing the distribution of Substate 4 neurons in WT and AD samples. Right: Chord diagram illustrating enriched biological processes and their associated genes in Substate 4 neurons. Links are defined as in **a**. **c.** Left: Scatter plot showing the distribution of Substate 5 neurons in WT and AD samples. Right: Chord diagram illustrating enriched biological processes and their associated genes in Substate 5 neurons. Links are defined as in **a**.

**Supplementary Note 8.** Enriched genes and pathways in hippocampal neuronal substates in MERSCOPE.

Beyond the isocortex, we extended our analysis framework to hippocampal neurons in the MERSCOPE dataset to investigate whether any substates exhibited disease-associated alterations. Using a similar pipeline, we clustered granule context embeddings and identified four hippocampal neuronal substates (Substates 1–4) shared between WT and AD samples (**Fig. S8a**). Each substate was defined by a unique composition of granule subtypes within its local microenvironment, as shown in the heatmap in **Fig. S8b**. Consistent with our previous findings, the Alluvial plot in **Fig. S8c** revealed little correspondence between these context-defined substates and those derived from somatic gene expression (ARI: 0.065), confirming that granule-based substates could not be identified through somatic transcriptome analysis alone.

We then asked how alterations in neuronal substates reflect the reshaping of neural networks under AD pathology. All substates showed distinct density distributions between WT and AD samples in the hippocampus. Most notably, Substate 1 neurons showed a significant increase in AD (**Fig. S8d, left**; two-sample t-test, p < 2.2 × 10⁻¹⁶) and were enriched for surrounding post-synaptic, dendritic, and ambiguously classified granules. As shown in **Fig. S8d (right)**, the enrichment of pathways related to dendrite development, protein localization to cell junctions, vesicle localization, membrane depolarization, and glutamatergic transmission suggests that this substate likely represents a population involved in postsynaptic integration, dendritic remodeling, and excitatory signal responsiveness.

In contrast, Substate 2 neurons exhibited a significant decline in AD (**Fig. S8e, left**; two-sample t-test, p < 2.2 × 10⁻¹⁶) and were enriched for surrounding pre-synaptic and ambiguously classified granules. As shown in **Fig. S8e (right)**, this substate showed upregulation of gene programs involved in SNARE complex assembly, synaptic vesicle cycling, regulation of calcium-dependent exocytosis, and amine transport, indicating a strong presynaptic identity optimized for regulated secretion and fast excitatory communication.

Together, the reduction of presynaptic-associated neurons (Substate 2) and the concurrent expansion of postsynaptic/dendrite-associated neurons (Substate 1) point to a critical presynaptic-to-postsynaptic rebalancing under early AD pathology, which destabilizes hippocampal circuits and ultimately drives the neural network toward hyperexcitability. This finding is consistent with our observations in the cortical neuronal population.

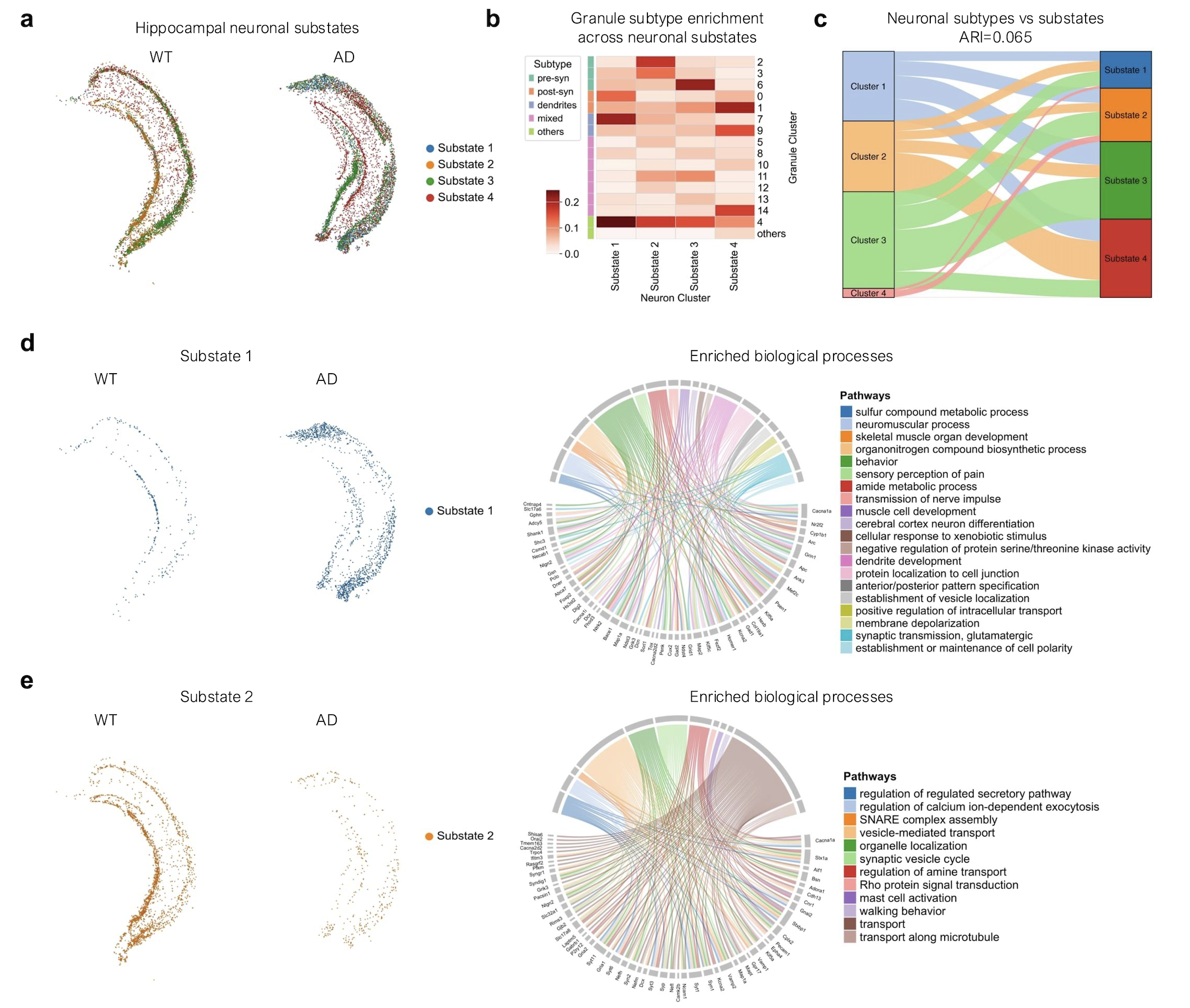

**Fig. S8. a.** Scatter plot showing the four hippocampal neuronal substates defined based on granule context. **b.** Heatmap showing the enrichment scores of granule clusters and their associated subtypes within each neuronal substate, calculated by averaging the granule context embeddings of individual neurons. **c.** Alluvial plot illustrating the correspondence between neuronal subtypes derived from somatic transcriptome clustering (left; K-Means clustering) and neuronal substates defined by granule context (right). **d.** Left: Scatter plot showing the distribution of Substate 1 neurons in WT and AD samples. Right: Chord diagram illustrating enriched biological processes and their associated genes in Substate 1 neurons. Links represent gene–pathway associations based on GO enrichment analysis. **e.** Left: Scatter plot showing the distribution of Substate 2 neurons in WT and AD samples. Right: Chord diagram illustrating enriched biological processes and their associated genes in Substate 2 neurons. Links are defined as in **d**.

**Supplementary Note 9.** Removing top-expressed markers in RNA granule detection in Xenium 5K.

To assess the robustness of mcDETECT in the absence of strong markers, we benchmarked the spatial granule distributions obtained from the full marker list against those derived after removing *Snap25* or *Slc17a7*, two of the top-expressed genes in the Xenium 5K dataset. As shown in **Fig. S9**, exclusion of either marker had little effect: regional granule densities remained highly correlated with the full-marker results, with Pearson correlations of 0.98 and 0.99, respectively. This robustness arises because mcDETECT leverages colocalization patterns across multiple granule markers, allowing it to infer underlying granule locations even when individual strong markers are absent.

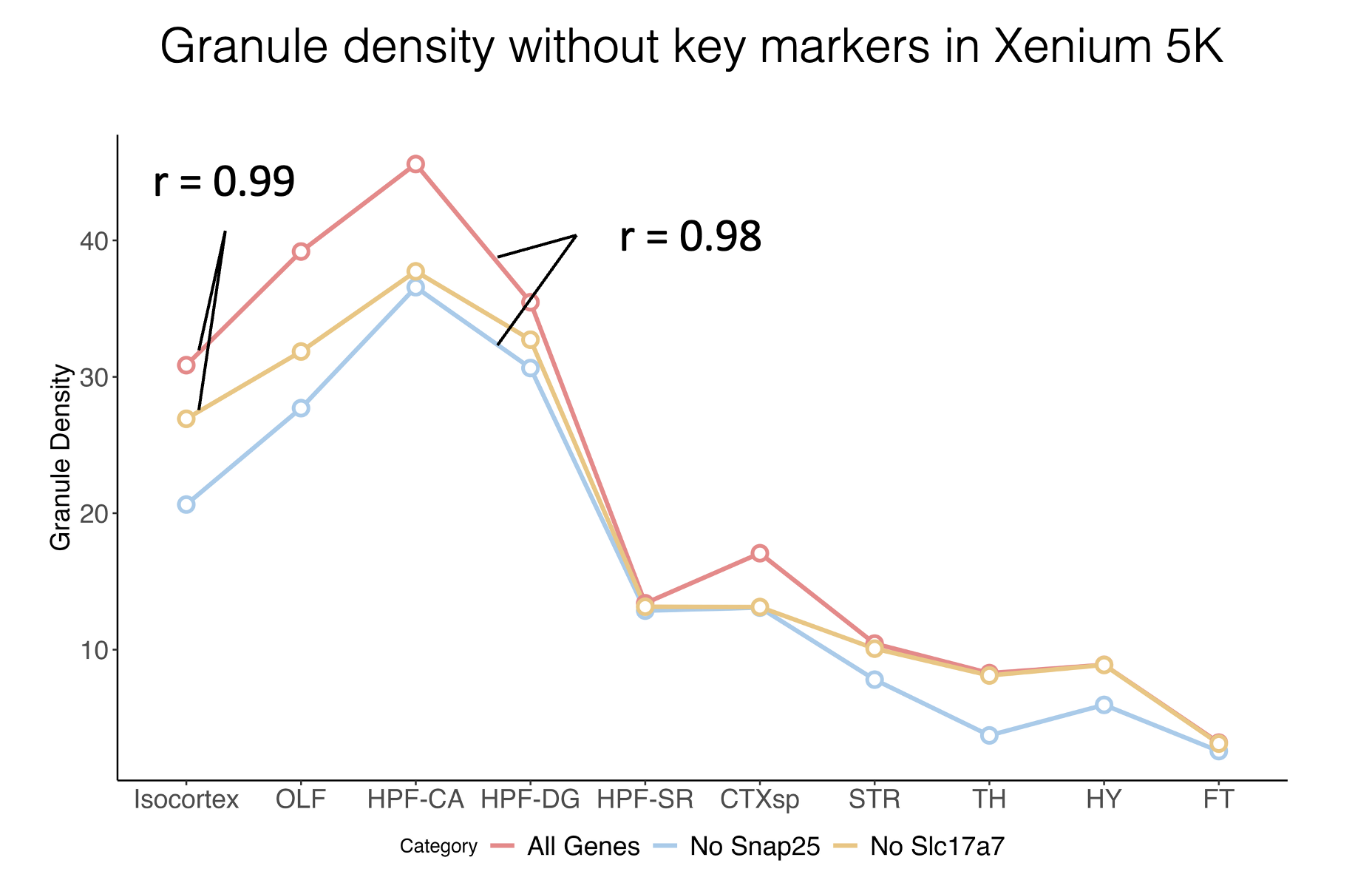

**Fig. S9.** Line plots comparing regional granule density trends in the Xenium 5K dataset using the full granule marker list, with *Snap25* excluded, and with *Slc17a7* excluded.
