## Supplementary Tables 5 and 6 for "Uncovering the dark transcriptome in polarized neuronal compartments with mcDETECT"

**Supplementary Table 5: Customized gene panel in the MERSCOPE experiment**

| | |
|---|---|
| 2010300C02Rik | |
| Abca7 | |
| Acsbg1 | |
| Adamtsl1 | |
| Adcy1 | |
| Adcy5 | |
| Adgrl4 | |
| Adora1 | |
| Aldh1a2 | |
| Aldh1l1 | |
| Ank3 | |
| Apc | |
| Apod | |
| Apoe | sequential panel |
| Aqp4 | |
| Arc | |
| Arhgap25 | |
| Arhgap6 | |
| B2m | |
| Bace1 | |
| Bcl11b | |
| Bdnf | |
| Bhlhe22 | |
| Bsn | |
| C1qa | |
| C1qb | |
| C1qc | |
| C4a | |
| C4b | |
| Cabp7 | |
| Cacna1a | |
| Cacna1i | |
| Cacna2d2 | |
| Calb1 | |
| Calb2 | |
| Camk2a | |
| Camk2b | |
| Camkk2 | |
| Cbln1 | |
| Cd11b | |
| Cd45 | |
| Cd63 | |
| Cdh13 | |
| Cdh6 | |

Chd9
Cldn5
Cnp
Cnr1
Cntnap4
Col19a1
Col1a1
Col6a1
Cpeb1
Cplx2
Cplx3
Cpne4
Cpne6
Cryab
Csf1r
Csmd1
Cspg4
Ctnna2
Ctss
Cux2
Cx30
Cx3cr1
Cyfip1
Cyfip2
Cyp1b1
Dcn
Dcx
Ddn
Deptor
Dgkz
Dkk3
Dlg1
Dlg2
Dlg3
Dlg4
Dlgap1
Dlgap4
Dner
Eno2
Epb41l1
Epha4
Fezf2
Fhod3
Fibcd1
Fign

Fmod
Fn1
Foxp2
Gabra2
Gabrb1
Gad1
Gad2
Gap43
Garnl3
Gatm
Gfap
Gfra2
Gja1
Gjb2
Gjc3
Gm2115
Gnai2
Gng12
Gphn
Gpr17
Gria1
Gria2
Grid1
Grik3
Grik5
Grin1
Grin2b
Grn
Gsn
Gucy1a2
Hexb
Hfe
Hivep3
Homer1
Homer2
Hpcal1
Hpcal4
Hs3st2
Htr1f
Iba1
Id2
Ifitm3
Igf2
Igfbp6
Inpp4b

Itgb4
Itpr1
Kalrn
Kcna2
Kcnh5
Kcnip4
Kif5a
Kif5c
Kmt2d
Ksr2
Lamp5
Laptm5
Ly6a
Map1a
Map2
Mapk4
Mapt
Mef2c
Meis2
Mog
Mpo
Myl4
Nav1
Ncam
Ndst3
Ndst4
Necab1
Necab2
Nefh
Nefl
Nefm
Nell1
Neun
Neurod1
Neurod6
Nfasc
Nlgn1
Nlgn2
Nlgn3
Nmral1
Nopchap1
Npnt
Npy2r
Nr2f2
Nrep

Nrg3
Nrn1
Nrxn1
Ntng1
Ntrk2
Nts
Ntsr2
Nwd2
Nxph3
Olig1
Olig2
Opalin
Or10ad1
Orai2
P2ry12
Pacsin1
Parm1
Pclo
Pde7b
Pdgfra
Pdyn
Pdzrn3
Pecam1
Penk
Pfkm
Plcxd2
Plcxd3
Plekhb1
Pou3f1
Ppp1r1b
Ppp1r9b
Prdm8
Prdx6
Prox1
Prph
Psd
Psen1
Psen2
Pum2
Purg
Pvalb
Rasgrf2
Rasl10a
Reep3
Rgs5

| | |
|---|---|
| Rims1 | |
| Rims3 | |
| Rmst | |
| Rnf152 | |
| Rorb | |
| Rprm | |
| Rspo1 | |
| Rspo2 | |
| Rtn4 | |
| S100a6 | |
| Satb2 | |
| Scg3 | |
| Shank1 | |
| Shank3 | |
| Shc3 | |
| Shisa6 | |
| Siglech | |
| Sipa1l1 | |
| Slc13a4 | |
| Slc17a6 | |
| Slc17a7 | |
| Slc17a8 | |
| Slc32a1 | |
| Slc39a12 | |
| Slit2 | |
| Snap25 | sequential panel |
| Sorcs3 | |
| Sorl1 | |
| Sox10 | |
| Spp1 | |
| Sptbn2 | |
| Stx1a | |
| Stxbp1 | |
| Sv2a | |
| Syn1 | |
| Syn2 | |
| Syndig1 | |
| Syngap1 | |
| Syngr1 | |
| Syp | |
| Syt1 | |
| Syt11 | |
| Syt17 | |
| Syt2 | |
| Syt3 | |

Syt4
Syt6
Tanc1
Thsd7a
Tle4
Tmem119
Tmem132d
Tmem163
Tmem255a
Tnf
Tox
Trpc4
Tubb3
Ube3a
Vamp1
Vamp2
Vat1l
Vcan
Vim
Vwc2l
Wfs1
Zbtb20
Zfpm2

**Supplementary Table 6: Long list of negative control markers**

Gm42617
Gm37249
Phxr4
A430110L20Rik
Gm43598
Gm10287
Tm6sf2
Gm44724
Olfr61
Gm13052
1700109H08Rik
Olfr46
Gdpd3
Phf2os1
Gm4258
Gm43174
Gm5577
Alkbh3
Gm26652
Muc3a
Neat1
Slc26a10
2700097O09Rik
6430590A07Rik
Robo3
Gm43279
Gm21781
Tnfrsf25
Cfp
Slc39a2
4930431P19Rik
Snhg20
Sec1
Gm14286
Gm15624
Tpcn2
Gm45205
Sfxn2
Syne4
Gm44806
Adam8
Cfap100
Gm4221
Xist

Gm43176
Ccdc134
Gm26901
Dleu2
Ccdc163
Gm44698
Slc2a4rg-ps
Gm17203
Miat
Mir337
B830012L14Rik
Mirg
Gm45178
Cd46
1700123O21Rik
Gm37899
Tmco6
Mroh7
Pdzd7
Gm37320
Gm15328
Mir124a-1hg
Kcnq1ot1
B230206H07Rik
Acrbp
9530059O14Rik
Gm37509
Sec14l5
Gm12216
Gm42878
Gm45179
Ftx
A330023F24Rik
Gm37238
Rgs11
Mir665
2610037D02Rik
Cdr1os
Gm3764
Arhgef1
Ppox
Apex2
Gm9930
Srpk3
Plekhn1

Col5a1
Ccdc84
Gm22373
Gm43175
Itga10
Gm28175
Il18bp
Zfp783
Firre
Gm31152
4930432B10Rik
4930594M22Rik
Tmc4
A230103L15Rik
Tia1
Gm45159
Adamts16
Thpo
4632427E13Rik
Fbln2
Miip
Stac3
Malat1
Lncpint
Acad10
Spaca6
Gm44257
4930539J05Rik
Gm38413
Arntl2
Adamts10
Gm42616
Ddb2
Gm3294
1700047M11Rik
Scarna2
Gm14636
4933439C10Rik
Gm12940
Gm9801
Acp6
Gm22205
Trmt44
Gm27032
2900005J15Rik

Gm27003
Arhgap17
Snhg17
Trmt13
Pask
Gm20045
Adamts20
Ankrd16
Mir99ahg
Lrrc29
Snhg11
BC065397
Gm26871
Olfr1564
Metap1d
Hps1
Tmem150a
Mamdc4
4732440D04Rik
Acaa1a
Rps6kb2
Tdrd5
Rbm3
1700012D14Rik
Gm44560
Gm19744
Flt3
Ints6l
Zkscan3
Leng8
2810428J06Rik
Rmdn1
Morn1
Zbtb49
Zcchc7
Col20a1
Tarbp2
Clasrp
Tyms
Chaserr
Gm16008
Jpx
Plcg2
Clcn1
Gm43327

Fbxl12
Ssh3
Gm43606
Gm4673
Gm37296
Crip3
Coro6
Wsb1
Gm12743
Odf2l
Snrnp70
Shkbp1
Cys1
Setd4
Cenpa
Safb2
Pdlim7
Mccc1
Helq
Stx3
Uckl1
Gm45250
Gcfc2
Cep57l1
Nemp2
Gm44997
Gm45869
Rskr
Ap1g2
Mri1
Nup93
Chrd
Tmem86b
Alg6
Gm37621
Hdhd5
Rreb1
Ccnl2
Hdac7
Gm37090
Gm27019
Amdhd2
Rnf207
A330076H08Rik
Snap23

Gm20754
Zgrf1
Col11a1
Meg3
Dennd6b
Tra2a
Wnk4
Tbce
Gm14443
Mtg2
C130073E24Rik
Atp6v0e
B230307C23Rik
Vmn2r85
Ccl25
A530013C23Rik
5730480H06Rik
Myo19
Fam228a
Ybx2
Gm42548
Prkn
Dmpk
Mthfs
Lmf1
Gm26703
Gm26621
Nsmaf
Cenpo
Zfp692
Caprin2
Usp40
Rnpc3
4930447C04Rik
Mthfsl
Nfkbid
Fam193b
9430025C20Rik
9330162G02Rik
P3h3
Hdac10
Cars2
Amn1
Tle2
Zfp950

Gm38073
Cfap65
Msantd2
Gm17334
Polg2
Chka
Ryr1
Tnip2
Prkd2
Gm14827
Hemk1
5830408C22Rik
Mtm1
Mov10
Sfi1
Thbs3
Clk1
P4ha2
Tmem128
Zc3h7a
Gm37928
Aifm3
3110021N24Rik
Cfap46
Nsun5
Tmem39a
Gm44136
E230016M11Rik
Sirt4
Dguok
Eif2d
B3gntl1
Flna
Man2c1
Neil1
Trim11
1110018N20Rik
5730522E02Rik
Ctnnal1
Krba1
Mir100hg
Dnajc24
Gm33533
Gm17494
Dvl2

Accs
Phykpl
Tbc1d2
Gm42549
Osgin2
Rassf1
Rhbdf1
Ubxn11
Mettl17
Slc35a3
Gm43379
Rbm5
Abtb1
Lrrc45
Rsrp1
Crlf1
A230077H06Rik
Poglut2
Cep95
Wdr90
A930012L18Rik
Rgl2
Clk4
Letm2
5031425E22Rik
Farsa
Anxa11
Pus7
St7
Junos
Abhd18
Czib
Cfap44
Stxbp2
Cyp4f13
Timm44
Gpt
Thumpd3
Stk38
Dcdc2b
Dusp11
Bmp1
Srsf11
Gm35339
Fam227a

Dgat1
Bbof1
Xkr6
Itga8
Plod2
Myo9b
Pdss1
Ttc14
Zgpat
Edem2
Coq6
Coq8b
Ttc21b
Cdk6
Il15ra
Ccdc28b
Dot1l
Ccdc159
Rccd1
Zfand1
Pan2
Tarbp1
Pot1b
BC030343
Sclt1
Kmt5c
Trank1
Plgrkt
Gm44559
Catsper2
Cdc7
Rpf1
Iqce
E230029C05Rik
Hectd2
Aspa
Slc16a11
Gm32444
Mlxipl
Khdrbs2
Cspp1
Gm44777
A930007I19Rik
Ermard
Prmt3

5530601H04Rik
Tmem181b-ps
Tatdn3
Slc38a6
Cep44
Arid3b
5430405H02Rik
Fbn2
Gm44033
Atg16l2
Vrk1
Lmbr1l
Zbtb8os
Pmm2
Fam53a
Rpia
Frmd4b
Igsf9
Anks6
Hook2
Gm30382
2810429I04Rik
Chuk
Matn4
Mlh1
Rdh13
Runx2
Josd2
Il1r1
Sfswap
Mecr
Tnxb
Rmc1
Ttll3
Pnpla3
Gm34294
Fam118b
Dus3l
Ankzf1
Pdia5
Col16a1
Telo2
9430064I24Rik
Dnah2
Tmem120b

5330434G04Rik
L3mbtl1
Col4a1
Tbc1d31
Adamts17
B9d1
Flnb
Pnkp
Bbs5
Gm44562
Recql5
Gtpbp6
Gtf2h3
Diablo
Slc16a10
Nfs1
Atad3a
Arglu1
Gmip
Nbeal2
1300002E11Rik
Pkn1
Fubp1
Anks3
Plxnb3
Rnft1
Traip
Osgep
Hexdc
2900052L18Rik
Pkd1l3
Abcc10
Smpd4
4732463B04Rik
Gm42756
Polb
Pop1
Abcd4
Actr5
Dnhd1
Stk19
Terf1
Slc26a6
Tamm41
Krit1

4732471J01Rik
4930570G19Rik
Gm29666
Mrgbp
Fam228b
Nfxl1
Trip6
Ccdc57
Il1rap
5430416N02Rik
Mcoln1
Iffo1
Strada
Rab34
Clcn2
Cttnbp2
Stx5a
1110038B12Rik
Swt1
Glt8d1
Ogt
Dnah9
Vwa3a
Gm37226
Gm44053
Snapc4
Dimt1
Ints13
Phkg1
Sgf29
Pxn
Zmym6
Hapln2
4833420G17Rik
Gm45605
P3h1
Malt1
Aox1
Ica1
Col27a1
Ppil3
Ssbp1
Luc7l2
Plk5
Cep78

Gtf2h2
Slc22a5
Ints10
Ap4e1
Bnip1
Tctn1
Fnbp4
Ulk4
Dpysl4
Sfpq
Lrrfip1
Rbm6
Nphp3
Reck
Wdhd1
Vwa3b
Rtel1
Gm5148
Egfem1
Akap8l
Zfp207
Dcp1b
Nxf1
Ikbkb
Rmnd1
Hdac8
Mfsd8
Tcte2
Tyw5
Arnt
Nktr
Mid1
Tmem67
Cep89
Col6a1
Cercam
Aasdh
Rbm39
Idua
Cfap74
Pus10
Exosc8
Med12
Cntrl
Taf6l

Acot8
Abhd14b
Taz
Chkb
Clk2
Itga7
Skp2
2810029C07Rik
Pnisr
Dhtkd1
Zfp446
Cemip
Ssc5d
Cdc25b
Sdccag8
Hps5
Cecr2
Shld1
Apobec3
Cdk9
Dock10
Gabpb2
Vmn2r87
Nsun7
Mapk1ip1
Cpsf4
Csf2ra
Ap5z1
Zfp57
Eml5
Pvt1
4930590J08Rik
Adck5
Stard9
Gmppa
Mrps10
Atxn7l2
A330102I10Rik
Ints8
Emx2os
Kptn
Tjap1
Lrrc1
Gsap
Gm13563

Usp16
Dtx2
E4f1
Pstk
1700016P03Rik
Gmcl1
Pvr
5031434O11Rik
A230057D06Rik
Slc15a4
Itga11
Mms19
Ranbp17
Hnrnpr
Ndor1
Carmil3
Zfp579
Hif3a
Nfkb1
Nat10
Ntn5
Mpzl1
Wdr54
Klhl12
Rpain
Casp2
Cep63
Orc4
Rcor3
Wrap73
Tfeb
Txlng
Sned1
Stx2
Ppp1r14a
Ccser1
Abcc8
Cep70
Celsr3
Dazap1
Mrps9
Hsf1
Ttc13
Hibch
Actl6a

Nrg2
Slc10a7
Rnaseh2b
Tulp3
4930503L19Rik
Pigz
Tmem208
Odf2
Hnrnpa2b1
Chchd7
Nphp4
Rras2
Tubd1
Prdm15
Gfm2
Smtn
Snapc3
Ccdc77
Cep192
Hace1
Lacc1
Rrnad1
Pdss2
Guf1
Crtc2
Gm42744
P2rx4
Ccl27a
Trim39
Ttll4
Tcea2
Tada1
Ankrd10
Gfpt2
Ddx55
Luc7l3
Gm38393
Mzf1
Smarcad1
Zfp945
Naa40
Abcc5
Prpsap1
Qtrt2
Prpf39

Plekhg5
2700049A03Rik
Paxbp1
Gprasp1
Mettl3
Ccdc30
Rbm25
Fxyd5
Tmem131l
Serinc2
Ctns
Nvl
Uggt2
Prmt9
Pcgf6
Pisd-ps1
Tchp
Anxa4
Focad
Slc50a1
Gm33989
Fus
Gripap1
Sfxn4
Ddx39b
Prpf40b
Gm37962
Pkd1
Usp48
Invs
Ltk
Gm47283
Zscan20
Tcirg1
Tars2
Sbno2
Rsu1
Aopep
Stk26
Cdan1
Ptbp1
Syn3
Cr1l
Rnf25
Dgcr8

Ryr3
Yeats2
Zcchc4
Ercc1
A230072C01Rik
Gtf2a2
Rfx1
Sirt7
Rrp9
Dph5
Best3
Mfsd10
Rabep2
Cdon
Ppih
Aldh1l2
Adgra2
Slc27a3
Rgs9
Fpgs
Dclre1c
Ltbp4
Med27
Traf7
Hnrnph1
Tmem29
Ints2
Rttn
Coa8
Mbd6
Trmu
Fuz
Hnrnpll
Fan1
D930048N14Rik
Med8
Phka2
Mettl8
Zfp61
Psme4
Gfus
Ilkap
Zfp672
Lrrtm4
Ccnl1

Rbm28
Cnppd1
Mterf4
Cenpv
Dhx35
Fbn1
Nfrkb
Vav3
Dcaf17
Snx29
Spns1
Nek10
Rapgef3
Adal
Mbip
Inpp5k
Rpusd3
Ifrd2
9430041J12Rik
Sirt6
Eif2ak4
Gm8066
U2af2
Ring1
Mtfmt
Trmt1
Mtmr10
Rab10os
Celf3
Rrp15
Pnpla6
Lonrf3
Tyk2
Paqr3
Flvcr1
Naa16
Slc26a11
Glmn
Pkp2
Prdm5
Abhd14a
Tango2
Dynlt2b
Taf1a
Thoc2l

Frmd4a
Dis3
Phldb1
Snx11
Dhx33
Mthfsd
Safb
Carns1
Anapc10
Tmem108
Engase
Tbcd
Dnttip1
Acss2
Top3b
Slc15a2
Ripk1
2810455O05Rik
Phf20l1
Srsf2
Tyw1
Pxdn
Rnf39
Col11a2
C130036L24Rik
Mbtd1
Tmem107
Ralgapa2
Bin3
Szt2
Plpbp
Akap8
Tmem87a
Ric1
Srek1
Gm16894
Dgka
Vwa5b2
St6galnac3
Pank4
Adat1
Lca5l
Fbxw9
Chfr
Nav2

Galnt11
Gga2
Hmces
Foxred1
Saal1
Smpd2
Pld1
Alpk1
Sec61a2
Ankrd24
Thap3
Irf3
Cpeb1
Map2k3
Setdb1
Slc12a9
Rbm10
E130307A14Rik
Dusp15
Srsf6
Rad9b
Trub1
Sec22a
Aloxe3
Gm43597
Slc2a6
Hjurp
Kctd18
Phkg2
Tmem240
Ece2
Grk4
Dus4l
Trmt2a
Eed
Khdc4
Stard5
Plpp5
Pgm2
2610005L07Rik
Dusp12
Slc25a40
Rrp1b
Gm10125
Ascc2

Tmem39b
Taf1c
Fubp3
Gm16638
Csad
Nup85
Dtymk
Lekr1
Grik2
Armc2
Trmt11
Psmf1
Mindy4
Slc66a1
Srrt
Tdp1
Alkbh1
Prkcd
Atp13a4
Selenoo
A830036E02Rik
A730060N03Rik
Hook1
Dock11
Srsf5
Snrpc
Nup35
Polm
Stxbp3
Lama5
Evc
Cnot10
Hdx
Dock6
Zc3h3
Dmap1
Zw10
Aig1
Tsen2
Gm6145
Supt20
Auh
Etfdh
Atp10b
Xrcc6

Mknk1
Tfr2
Dpp7
2010315B03Rik
Elovl1
Zfp263
Marchf7
Slc25a26
Sgk3
Iqgap1
Pex1
Chpf2
Zfyve16
Mpp3
Fahd2a
Utp25
Donson
Trabd
Nars2
Ss18
Arhgap39
Stk36
Slc25a42
Ppip5k2
Sntg1
Katna1
Frmd5
2300009A05Rik
Dnm2
Caap1
Atp11b
1600020E01Rik
Nrf1
Hyls1
Morc2b
Rbm22
Ddx27
Dph1
Dxo
Zfc3h1
Gm44644
Usp28
Malsu1
4933421O10Rik
Unk

Dop1a
Tmbim1
Ankrd54
Hps4
Gm43268
Plekha7
Zfp287
Fance
Supt3
Sft2d1
Mapk11
Eml6
Cdk8
Rab36
Sgcd
Prpf38b
Txnrd2
Ano4
Eln
Smarcd2
Ice2
Npr2
Ehmt1
Plekhh2
Nup160
Borcs8
Cwc22
Wdr83
Lig3
Ddx49
Pgls
Nudt13
Gmpr2
Tsen15
Cdk5rap1
Bicc1
Galnt3
Mindy3
Myt1
Inpp5b
Fam133b
Nrbp2
Camkmt
Exosc7
Cep43

Lrwd1
Adamtsl4
Ebf4
Pick1
Slc7a6
Ppfibp2
Pnpt1
Zfat
Lmna
Spns2
Ecm1
Haghl
Tbp
Wdr70
Dnajb4
Trmt1l
Trpc6
Gm38192
Polr3e
Snrnp48
Ypel4
Psmd9
Gm16835
Zfp653
Nup205
Cox18
Exoc1
Slc66a2
Pik3c2b
Orc6
Ccdc174
Acad8
Gigyf1
Shfl
Heatr6
Rnpepl1
Phaf1
Chic2
Inha
Arrdc1
Lrif1
Obsl1
Cop1
Scarb1
Piga

Mkx
Rgsl1
Plag1
Iqcc
Appl2
Crlf3
C2cd5
N4bp2l2
Slc26a4
Sox2ot
Crebzf
Katnip
Ssr4
Plekhg3
Nsun4
Cfap54
Rbm33
Thoc1
Cars
Tm6sf1
Xylb
Pld2
Trit1
Dph7
Klhdc4
Plcd1
Stat2
Fancc
Ftsj1
Wrap53
Rhno1
Ppil2
Crocc
Myg1
N4bp2
Fam76b
Cdk5rap2
Itfg2
Nemp1
Whamm
Pyroxd1
Cep112
Epha10
Alg13
Trnau1ap

Golga1
Nubp1
Mdm4
Rnpep
Cabp1
Stx18
Zfp169
Teddm2
Ppcdc
0610030E20Rik
Ccnt2
Prtg
Adprs
Rufy2
Zfp280d
Cntrob
Katnal2
Wdr20
Heatr3
Ift172
Atg16l1
Osbpl7
Tgds
Xrcc1
Slc16a13
C030029H02Rik
Agk
Faap100
Fn3k
Zfp386
2900076A07Rik
Kcnn2
Mrps35
Col4a2
Las1l
Akr1b3
Ggcx
Fam229b
Zfp157
Grb14
Marchf11
Ttc4
Slc26a8
Fbxo27
9930104L06Rik

Ptcd3
Suds3
Adam17
Armc9
Ddx31
Ewsr1
Pwwp3a
Apeh
9530026P05Rik
Prpf38a
Dlk2
6430584L05Rik
Zfr2
Ugdh
Agrn
Carmil1
Pcyt2
Slc9a8
Prkd1
Uhrf2
Clk3
Nae1
Msra
Mthfd2l
Ighm
Jakmip3
Gmds
Csf3r
Pigu
Ift140
Gm2824
Slc24a4
Trp53
Myef2
Pnpla7
Pisd-ps2
Trafd1
Dnajc1
Kcnh4
Senp1
Plscr3
Cnot3
Slc25a36
Shc1
Prr5l

Cplane1
Rwdd3
Acadvl
Arv1
Elmod2
Atf1
9430015G10Rik
Phf21b
Acad9
Ncapd3
Pan3
Tctn3
Tmem38b
Fnbp1
B4galt1
Tmem145
Sat2
Aste1
Cyfip1
Pms2
Smim20
Crtc3
Cfap69
Glb1l
Zcchc9
Zswim9
Zfp182
Tra2b
Bud23
Bcat2
Prodh
Chid1
Car7
Pign
Emc9
Nrg3
Gss
Rabl2
Hnrnpm
Lrch1
Acot9
Ccdc39
Slc25a35
Pcgf2
Rarg

Nhlrc3
Atp10a
Tial1
Mettl5
Rbm41
Mterf3
Tex9
Fbxl6
Ppan
Ppp4r1l-ps
Cog2
Cc2d2a
Mok
Snhg14
Zkscan2
Sgpl1
Pced1b
Tent2
Aifm1
Pop4
Sec11a
Sntb2
Plod3
Cers4
Mier2
Cdh24
2210408F21Rik
Tcf3
Gm15446
Ift122
Tle1
Ercc2
Rars2
Agbl5
Rbmx
Rnf121
Fbxw4
Rnf32
Morc4
Stard13
Atr
Meis3
Sgce
Arap1
Fam214a

Snrnp40
Tbc1d19
Pcolce2
Cc2d1b
Serac1
Cpne6
Sbf2
Fancg
Dzip1l
Ddx47
Pcsk7
Porcn
Gnl2
4933427D14Rik
Zfp408
Dus2
Epb41
Spice1
Slx1b
Dennd3
Vezt
Svil
Spata7
B3galnt2
Zfp101
Zdhhc1
Mus81
Il11ra1
Izumo4
Atp13a1
Atxn2l
Spag1
Ip6k2
Sppl2b
Pusl1
Jmjd6
Rbck1
Stra6
Slc29a4
Pced1a
Snx32
Senp7
Prpf18
Agbl3
Lbr

Ubp1
Rfc5
Sat1
Top3a
Trdmt1
Nup88
Nelfa
Pcdh15
Map3k3
Ttc8
Gtpbp2
Frmd8
Mfsd11
B130024G19Rik
Pigv
Cep135
Aagab
Ippk
Vav2
Efcab6
Itpkc
Slc2a8
Utp14a
Ercc8
Polr3gl
Nup188
Pde7a
Tubgcp4
Nup210
Irf9
Arrb2
Pkd2
Raf1
Atg3
Ybx3
Slco1c1
Col6a2
Dock7
Srsf9
Nfkbiz
Mrpl23
St8sia4
Tex10
Ganc
Ttc37

Map2k5
Scnn1a
Mrps25
Arfrp1
Prpf3
Map3k14
Zfp335
Tet2
Tma16
Zfp330
Fam219b
Usp19
Fbl
Susd1
Dhx16
Fam184b
Mocs1
Hbp1
Gm17354
Gin1
Arhgef10
Nr2c1
Sugp2
Mcf2l
Gm5141
Hspa14
Gls2
Tsga10
Slf2
Scaf4
Arhgef40
Uvssa
Eftud2
Smg9
Zcchc8
Eif2b3
Tspan18
Hps3
Unc13b
Ggt7
Kmt5b
Trpm7
Rae1
Slc39a13
Plekha1

Brat1
Hars2
Gm2762
Vps33b
Map4k5
Brwd1
Tbc1d17
Ddx41
Cfap410
Adsl
Hebp1
Gm10516
Kif17
Rpgr
Htr4
Cers5
Cacna1d
Cpne9
Tardbp
Dgkq
Ppp4r1
Samd5
Rffl
Orc3
Rnf149
Arntl
Pigb
Gpr107
Ttc39c
St7l
Cdc23
Rilpl2
Onecut2
Zfp384
Cln3
Prr14
Txndc11
Kmt2a
Denn2b
Zfp280c
Lama3
Rbm12b2
Jag1
Coro7
Kdm4c

C4a
Lrrc42
Cep162
Ptpn2
Rev1
Lpcat3
Rian
Nle1
Sema5b
Eml1
Pdrg1
Stard3nl
Nprl3
B230369F24Rik
Hadhb
Slc29a1
Me3
Abhd1
Plxna3
Gdpd5
Rogdi
Mdn1
Thap4
Eps15l1
Rfc2
Sun1
Usp54
Rxfp1
Slc35f4
Col19a1
Dnajb5
Osbpl3
Tspoap1
Exosc5
Pla2g4e
Cybc1
Tpra1
Pskh1
St18
Anks1
Zmat4
Adamts9
Card10
Cacna1h
Rex1bd

Alg12
Apba3
Bbs7
Ptchd4
Tmem161b
Aatf
Slc29a2
Cdk4
Fbxo8
Pde9a
Sra1
Jmjd1c
Polrmt
Plat
Zbtb40
Dis3l2
Mir9-3hg
Prim2
Zbtb37
A330008L17Rik
Zfp639
Cpsf6
Kdm1b
Ccdc130
Srsf3
Nrip2
Zranb2
Ddx39a
Srrm1
Rnf112
Dock5
Aspscr1
Glis3
Actl6b
Sema4c
Dcbld1
Atp11c
Lrrcc1
Pold3
Sem1
Prkag2
Fnip2
Cibar1
Lman2l
Slc25a27

Plekha2
Mcph1
Mettl15
Psrc1
Dgki
Plekha5
Anapc5
Zfp598
Trpm4
Antkmt
Cald1
Man2b1
Tppp3
A830073O21Rik
Tubgcp3
Cep97
Slc35f6
Sptbn4
Ncapd2
Tnrc6a
4921531C22Rik
Timm23
Idh2
Csmd2
Rad9a
Gdpd2
Lrrfip2
Ifi27
Chd7
Shtn1
Acin1
Pcca
Clvs2
Surf2
Ncbp3
Plbd2
Rft1
Git2
Zfp112
Pard3
Rnf31
Inpp5j
Gm15506
Myo1c
Ears2

Zbtb48
Polg
Pomt1
Opalin
Uimc1
Lct
Inpp5e
Tmem198b
Zfp964
Lrch3
Syt14
Igsf9b
Dnaaf9
Cyb5r4
Gale
Lnpk
C130071C03Rik
Ciz1
Gps2
Wdr19
Zfp746
Brd8
Kif13b
Alkbh4
Tubgcp2
Zxdc
Brd9
Vipr1
Prkrip1
Lipe
Ash2l
Pomt2
Csmd3
Mfsd2a
Phip
Ciart
Htra2
Mios
Hinfp
Sidt1
Sec24a
Lpcat2
Carf
Lemd2
Ryk

D11Wsu47e
Kctd6
Smarcb1
Slc31a2
Eef2k
Usp36
Pofut1
Gm43457
Ttf1
Srrm2
Rsrc2
Cep164
Pkn2
Smg1
Emg1
Lsm8
Chd1l
Cdkal1
Gtpbp3
Sptlc1
Dus1l
Cd2ap
Rbm26
Polr3f
Frg1
Rdh10
Lmnb2
Arid5a
Nbeal1
Fam98c
Tlcd3a
Gpr108
Dpf2
Armc6
Trpc4
Dync1i2
C1ql2
Gba2
Raver1
Matk
Ntmt1
Tab1
Naca
Spag4
Gt(ROSA)26Sor

Vcl
Atg4b
Enox1
Galns
Mtdh
Pou2f1
Riok1
Zscan22
Gm10649
Wdr75
Ppfia1
N4bp2l1
Dalrd3
Ppip5k1
Slit1
Naxd
Echs1
Gm7694
Btaf1
Nr6a1
Marveld1
Sh3gl1
Pmpcb
Rce1
Shc2
Dennd1b
Tcf12
Pex10
Ccdc66
Zfp174
Trub2
Lactb2
Macir
Ephb2
Bicd1
Cpsf1
Cops7b
Gm37607
Patj
Tti2
Cxxc1
Jam3
Sik2
Spata5
Nt5c3b

Mindy1
Rad52
Far1
Rchy1
Zfp329
Spag6l
Fam20a
Tor2a
Arfgap1
Nup54
Aamdc
Zfyve28
Hic2
Zfp512b
Acvr1
Mettl2
Clcc1
Riok3
Spag5
Ddx51
Fam3a
Nol11
Alkbh6
Adgrb3
Kcnt1
Csmd1
BC034090
Abca4
Kantr
Tnik
Slc35a1
Abca7
Dnai4
Stk10
Kdm3a
Aars2
Gabpb1
Srsf10
Washc4
Kctd16
Tmem214
Zfp583
Ikzf2
Polk
Slc9a9

Ccdc9
Pcsk5
Mavs
Slc7a10
Cog3
Zbtb1
Pou2f2
Mpi
Kri1
Hdac1
Fbxo6
Ccnjl
Garnl3
Prdm10
Ptdss2
Zfp324
Gpatch8
Lta4h
Lamtor3
Fam114a2
Tead1
Fah
Wdr3
Nfatc2
Bbs2
Thsd7b
Tmem80
Zfp235
Mtx1
Faim
Tmem134
Faf1
Mib2
Pigo
Mical2
Mdga2
Galc
Mia2
Dnaaf5
Skiv2l
Mthfr
Pde3b
Daglb
Ptpn23
Gm43682

Yipf1
Slc44a5
Stambp
Afap1
Mical3
Slc4a7
Ercc4
Mtf2
Hacd1
Scnm1
Phtf2
Taf15
Gtpbp8
Gramd2
Plekha6
Cdc14a
Sass6
Uevld
Med23
Ngly1
A930005H10Rik
Kmt2b
Ppp4c
P3h2
Pfas
Olfml2b
Radil
Cope
Lyst
Gpnmb
Fam117a
Ccdc120
Slc39a14
Slc22a15
Gjc3
Gba
Secisbp2
Prox1
Atpsckmt
Trmt6
Urb1
Wdr46
Mcam
Rnf215
Fuom

Gucd1
Atp11a
Thop1
Chsy3
Tmem219
Hdac3
Pot1a
Tubgcp5
Mcts1
Cttn
Ern1
Gga3
Zfp511
Wdr61
Xpo4
Mphosph9
Mettl22
F730043M19Rik
Spg11
Rps6kl1
Med24
Cacnb2
Npepl1
Rfxank
Mrpl24
Ptbp2
Tmem25
Yars2
Abhd11
H13
Arsj
Stab1
Eef1d
Tango6
Edrf1
Hspbap1
Usp4
Gk5
Dmtf1
Hdac6
Utp20
Mllt10
Commd4
Atxn7
Slc20a1

R3hdm1
Mtmr2
Katnbl1
Nek1
Ecpas
Ankrd27
Zfp420
Map4k2
Dtx3
Relch
Sh2b2
Cwc27
Glt8d2
Dclk2
Cox19
Cxcl12
Mroh1
Ablim2
Zfp326
Fbxo7
Cyth1
Tbc1d10a
Bnip2
Ccdc9b
Tank
Rngtt
Srpk1
Mpc1
Mrpl15
Sec13
Plekhh1
Thoc5
Dtnb
Eef1aknmt
Sec24b
Sh3glb2
Tspan33
Ift80
Rfx2
Dcp1a
Mrnip
Ago3
Pcgf1
Ago4
Mdc1

Birc2
Ttc17
Stambpl1
Abcc4
Taf1b
Mbd1
Zfp28
Cluap1
Ophn1
Sepsecs
Mtch2
Limk2
Utp11
Creb5
Cyth4
Bcas2
Mrps18b
Frem1
Ndufb7
Scfd2
Rcbtb2
Sf3b1
Nin
Msh3
Vps13c
Med6
Rhot2
Aspdh
Nmrk1
Katnb1
Adam15
Zfpl1
Paip2
Luc7l
Vps45
Tfcp2
Dnah1
Ttc7
Pibf1
Swi5
Kctd17
Ppif
Nfya
Gpsm2
Mccc2

Dhx32
Map3k4
Cad
Nup98
Kyat1
LTO1
Sgk1
Ivd
Gm4924
Ppp2r3d
Tm7sf2
Abcd1
R3hcc1l
Wdr66
Zfyve27
Ankrd13a
Tmem175
Numbl
Cep85l
Ppp1r12c
Lurap1l
Klhl32
Nfyc
Sorbs2
Slc27a1
BC002059
Sumf2
Map7
Shprh
Cdc26
Erbb3
Creld2
Tbata
Ptk7
Zmym3
Smarce1
Atp8b2
Baz1b
Drg2
Arih2
Pds5a
Mpc2
Lrig2
Fbrs
Grin2c

Ctps2
Dmxl1
Nasp
Vps35l
Ahctf1
Lamb1
Ino80e
Rsad1
Manba
Slc30a7
Gcdh
Tmem63a
Prkab1
Larp1b
Cenpc1
Ulk3
Nbas
Unkl
Atg4c
Baz2b
Elp1
Per2
Pias4
Slc37a1
Asxl3
Nol9
Adarb2
Gm21092
Emsy
Mob2
Psma3
Maip1
Mast4
G2e3
Cdkl1
Grip1
Stn1
Commd7
Cacna1g
Dpyd
Iws1
Zfp398
Nsfl1c
Taf1d
Tbc1d12

Eri2
Grhl1
Per1
Ttll5
Acap2
Abcc1
Mcmbp
Grik4
Fbxw2
Lsg1
Thada
Mrpl37
Elmod3
Unc45a
Dapk3
Rapgef6
Lratd2
Trpv2
Rbms1
Pla2g12a
Nrros
Alg5
Proser1
Ric8a
Mnat1
Sema4b
Tor1b
Gmeb2
Wtap
Ppa2
Grik1
Ap3m1
Cyb561d1
Ell
Kdm2b
Edc4
Mtmr1
Rbpj
Selenon
2610035D17Rik
Sdad1
Zfp667
Fibp
Gnl3
Vcan

Zfand2b
Fbf1
Fbxl5
Cenpt
Ankrd42
Lcor
Cep131
Cables1
Zfp944
Snx1
Ddx18
Dok7
Zfyve26
Washc3
Ccdc93
Rab3gap2
Pip4p1
F3
Msto1
Snx24
Prkx
Sema6c
Pwwp2b
Zbtb17
Gramd1a
Skap2
Taf2
Rnf20
Ppid
Gm37305
Mfap3
Psph
Lmtk3
Fndc1
Gtf2b
Zcrb1
Galk2
Rtkn
Nipal2
Arhgap18
Naa20
Blvra
Srrm3
Sars2
Mark3

Myh9
Septin11
Ext2
Polr2e
D5Ertd579e
Fchsd2
Fastkd1
Lrp1b
Tgfbr3
Jakmip1
Gpaa1
Pstpip2
Med17
Ahcyl2
Aph1a
Picalm
Ddx17
Slc38a9
Uvrag
Rbak
Son
Ift46
Adamts3
Nit1
E130308A19Rik
Arpc5l
Adh5
Cog1
2310061I04Rik
Eme2
8430429K09Rik
Polr1b
Pfkfb4
Ciao3
Ddhd1
Surf6
Iqcb1
Utp4
Unc79
Mat2a
Elapor1
Zfp2
Psmd6
4933434E20Rik
Abcf3

Akap13
Dnah6
Dgkd
Scyl3
Mak16
Nhlrc2
Abhd16a
Zfp560
Raver2
Galnt18
Ints6
Colgalt1
Cep295
Arpp21
Sorbs3
Npnt
Mtif3
Dync2i1
Bak1
Atp7a
Zfp111
Zfand6
Ubr5
Dhodh
Kansl2
Zmynd8
Nfx1
Kdm6a
Tpp1
Slc35d1
Mtx2
Polr3b
Gpld1
4632404H12Rik
Cstf3
1110002E22Rik
Epha7
Slc52a2
Ist1
Osbpl5
Tmpo
Exoc7
Lamc1
Napa
Stxbp6

Slc25a19
Asxl2
Tfdp2
Crot
Fndc3b
Ube2j2
Dpf1
Tpd52l2
Hnrnpdl
Lama2
Kcnj6
Cpt1c
Phf14
Dhx15
Mau2
Rbm4b
P2rx7
S100pbp
Plcg1
P4ha1
Vars2
Pola1
Rab28
Gtf3c5
Rab24
Rint1
Upf3b
Csnk1g1
Atp9b
Dcaf15
Bcan
Bicdl1
Zfp523
Aup1
Slc16a6
Agbl4
Tmod1
Ovca2
Hdgfl2
Pfkp
Lair1
Brdt
Ncoa3
9230112E08Rik
Fbrsl1

Cntn5
Ptcd2
Tada2a
Entr1
Cobll1
Scfd1
Dhx30
Lias
Spsb1
Trim3
Timm9
Chd8
Mmaa
Aqr
Mospd2
Tdg
Dcaf8
Klhdc8b
Alms1
Cfap20
Zswim8
Pxylp1
Noc3l
Mysm1
Heatr1
Shisa7
Rheb
Plcd4
Fyco1
Ndst3
Ampd2
Arl13b
9130401M01Rik
Nupl2
Anapc1
Stard3
Memo1
Zbtb25
Phf1
Ranbp10
Coq10b
Fkbp15
Dok6
Pex16
Eral1

Pex26
Dctn6
Zfp961
Sgms1
Snx14
Fam13a
Lrpprc
N6amt1
Brwd3
Kcnc3
Ap4m1
Pls1
Gpatch2
Suclg1
Uba5
Slc11a2
Samhd1
Ccdc137
Nsmce2
Zfhx4
Zfp426
Smchd1
Bcl11a
Gpr137b-ps
Slco5a1
Capn7
Nabp1
Prkg1
Zfp189
Srsf1
Mpp1
Fnta
Cyth3
Atad5
Rusf1
Pbx4
Klhl2
Zfpm2
Hnrnpl
Hnrnph3
Mier1
Qsox1
Snx15
Eogt
Gm12258

Fam149b
Akr7a5
Tmem165
Kat6b
Utp18
Ttc19
6820431F20Rik
Psmd13
Gcn1
Trmt61a
Atp6v1h
Calcoco1
Phtf1
Hykk
Rftn1
Cwf19l1
Vasp
Npdc1
Mir124-2hg
Cul9
Pdcd11
Aaas
Timm50
Ubiad1
Ndufab1
Zfp445
Pex3
Cdc14b
Gm45053
Itpa
Uspl1
Zfp282
Pik3ip1
Ccdc86
Gria2
Kl
Rnf123
Chd6
Stag1
Uros
Tmem201
Kcnq4
Ints9
9130024F11Rik
Sec11c

Gk
Mtg1
Chd2
Gpr162
Ache
Nckap5
Vps8
Ankrd44
Tmem62
Ankhd1
Slc30a6
Commd3
Vars
Sel1l3
Grm7
Il6ra
Dcun1d2
Trp53bp1
Acox3
Rps6kb1
Traf3ip1
Vps54
Gm43668
Mitf
Pgd
Klhl20
Usp21
Scmh1
Ttc26
Ube2d1
Ubn2
Rbbp4
Mybbp1a
Antxr1
Mmp16
Snta1
Spata13
Noc2l
Zfp948
Atat1
Arfip2
Kirrel3
Hbs1l
Rpap1
Mrpl58

Ints11
Dclre1a
Arhgef2
Haus6
Zfp532
Plekhm1
Tmem161a
Vrk3
Lrfn5
Slc37a4
Hmgcl
Idh3b
Parn
Tpp2
Xpo6
Kat8
Sharpin
Scamp2
Cep250
Wdr59
Rabggta
Ctdspl2
Efcc1
Wdr4
Slc6a6
Psmb3
Ppp2r2d
Derl2
Kndc1
Yif1a
Rbm45
Dpf3
Xab2
Idh3g
Zfp949
Pus3
Med25
Zfp691
Disp3
Clcn7
Kdm4b
Myrf
Brms1
Abcb8
Fam135a

Tiprl
Cabin1
Tbc1d23
Lca5
Ndst2
Tln1
Kremen1
Rnf2
Trmt10b
Ahsa2
Dgkh
Atg2a
Bms1
Gnptg
Necap2
Azin2
Anapc4
Nob1
Top1
Dennd6a
Mrpl13
Fbxo44
Baiap2
Capn15
Fam135b
Gnpda2
Cblb
Lsm1
Ralgps1
Nom1
Mvb12a
Tbc1d24
